# A hybrid high-resolution anatomical MRI atlas with sub-parcellation of cortical gyri using resting fMRI

**DOI:** 10.1101/2020.09.12.294322

**Authors:** Anand A. Joshi, Soyoung Choi, Yijun Liu, Minqi Chong, Gaurav Sonkar, Jorge Gonzalez-Martinez, Dileep Nair, Jessica L. Wisnowski, Justin P. Haldar, David W. Shattuck, Hanna Damasio, Richard M. Leahy

## Abstract

We present a new high-quality, single-subject atlas with sub-millimeter voxel resolution, high SNR, and excellent grey-white tissue contrast to resolve fine anatomical details. The atlas is labeled into two parcellation schemes: 1) the anatomical BCI-DNI atlas, which is manually labeled based on known morphological and anatomical features, and 2) the hybrid USCBrain atlas, which incorporates functional information to guide the sub-parcellation of cerebral cortex. In both cases, we provide consistent volumetric and cortical surface-based parcellation and labeling. The intended use of the atlas is as a reference template for structural coregistration and labeling of individual brains. A single-subject T1-weighted image was acquired at a resolution of 0.547mm **×** 0.547mm **×** 0.800mm five times and averaged. Images were processed by an expert neuroanatomist using semi-automated methods in BrainSuite to extract the brain, classify tissue-types, and render anatomical surfaces. Sixty-six cortical and 29 noncortical regions were manually labeled to generate the BCI-DNI atlas. The cortical regions were further sub-parcellated into 130 cortical regions based on multi-subject connectivity analysis using resting fMRI (rfMRI) data from the Human Connectome Project (HCP) database to produce the USCBrain atlas. In addition, we provide a delineation between sulcal valleys and gyral crowns, which offer an additional set of 26 sulcal subregions per hemisphere. Lastly, a probabilistic map is provided to give users a quantitative measure of reliability for each gyral subdivision. Utility of the atlas was assessed by computing adjusted Rand indices between individual sub-parcellations obtained through structural-only coregistration to the USCBrain atlas and sub-parcellations obtained directly from each subject’s resting fMRI data. Both atlas parcellations can be used with the BrainSuite, FreeSurfer, and FSL software packages.

## 1. Introduction

Atlas-based identification of neuroanatomy plays an integral role in studies of neurological disease, cognitive neuroscience, development, and aging, as well as in clinical applications including neurosurgical and radiation treatment planning (Dickie et al., 2017; Taylor et al., 2020). Utilizing a reference atlas, these approaches parcellate human brains into homologous brain regions for comparison of regional quantitative measures such as brain volume, cortical thickness, microstructural diffusion, blood perfusion, and functional activity. While atlases can be study-specific, publically available atlases are advantageous in that they allow for direct comparisons across studies and study populations. Many of the publically available human brain atlases have been derived from structural anatomical data. These rely almost exclusively on structural features, namely, sulci and gyri, to define homologous brain regions. However, studies have shown that anatomically-defined cortical brain regions are often heterogenous and comprise multiple subregions with distinct functions. However, the optimal number of subdivisions per cortical brain region remains unknown. Likewise, the degree to which consistency of subdivisions across individuals varies across brain regions is unknown. The novel atlas described below combines structural delineation of gyral anatomy with functionally-driven subdivisions of each gyrus. We assess the consistency of each subdivision across a group of independent subjects. This hybrid combination of structural and functionally determined regions may prove to be a useful addition to the tools available for quantitative brain analyses.

Modern imaging techniques have expanded the range of properties and the resolution with which we can observe them in the brain. Many alternative methods of brain parcellations have been explored to match the diversity and resolution of brain parcels to those of the observed findings. A range of imaging methods have been used to identify cytoarchitectonic, myeloarchitectonic, or functional features that are used in turn to parcellate the brain (Chakravarty et al., 2006; Chong et al., 2017; Essen and Drury, 1997; Yeo et al., 2011; Zilles and Amunts, 2010). White matter architecture has been used to parcellate the cortex based on differences in myelination within distinct brain regions (Glasser et al., 2014; Glasser and Van Essen, 2011), as well as through delineation of regions based on structural connectivity between these regions (Behrens et al., 2003; Johansen-Berg et al., 2004). Similarly, functional MRI (fMRI) can be used to identify contiguous regions of cortex that show similar functional properties (Bhushan et al., 2016; Joshi et al., 2018; Margulies et al., 2009). Parcellation methods based on fMRI include region growing (Blumensath et al., 2012), graph cut (Chong et al., 2017; Li et al., 2018; Shi and Malik, 2005), hierarchical clustering (Cordes et al., 2002), and ICA (Calhoun et al., 2009). Individual variations and limited data from single-subject templates may not be representative of the larger population. To address individual variation, multi-subject templates, including average brain and probabilistic atlases, attempt to account for the variability of brain architecture across a population (Desikan et al., 2006; Fischl, 2012; Mazziotta et al., 1995). Multi-subject functional atlases are created by anatomical coregistration onto a common space followed by group parcellation based on fMRI (Chong et al., 2017; Craddock et al., 2013; Glasser et al., 2016; Yeo et al., 2011). Lastly, multi-modal and multi-subject approaches have pooled data across subjects and with different imaging contrasts to build an atlas with an integrated pattern classification strategy (Glasser et al., 2016). Glasser and colleagues used a large number of multimodal features from Human Connectome Project (HCP) data to parcellate the cerebral cortex into 180 unique regions per hemisphere (Glasser et al., 2016). A significant effort was made to understand the relationship of this parcellation with other atlases (See supplementary results in (Glasser et al., 2016)), but the final parcellation is largely independent of sulcal and gyral boundaries and therefore continuity with earlier studies and literature is hard to establish.

It is important to note that the studies and atlases cited above have not agreed on a common parcellation and each method has its own assumptions (Zhi et al., 2021). Given the broad array of available techniques and atlases, selection of the most appropriate method of parcellation or choice of atlas depends heavily on the application. Arguments, however, can be made for the continued use of anatomical segmentations. For example, a practical challenge in using parcellations that are not based on standard anatomical landmarks is how to compare findings with earlier studies based on classically defined gyral labels. Regions in newly defined parcellations can overlap with multiple historically defined regions, leading to ambiguity in referencing these regions for meta-analysis. Continuity with anatomically-defined historical parcellations is also important, for example, in neurosurgery applications (Taylor et al., 2021). While surgical resections for treatment of epilepsy and brain tumors often use task-fMRI based identification of eloquent cortex, resection volumes are typically defined with respect to anatomical landmarks (Makris et al., 1999; Mori et al., 2013). A parcellation scheme that has consistency with anatomical labels based on well-known sulcal and gyral landmarks is therefore important in the context of surgical planning and research studies of outcomes.

While sulcal landmarks are the first salient anchors for cortical subdivisions, the associated gyral regions are often large relative to resection volumes. Linking gyral sub-parcellations to functional specialization based on fMRI connectivity provides a principled basis for defining smaller parcels. This in turn could provide surgeons information to better delineate focal resections and facilitate inter-subject comparison of outcomes. Similarly, mapping invasive stereotactic EEG (SEEG) recordings to the atlas (Taylor et al., 2021) allows for population studies of the relationship between epileptiform activity with respect to a consistent labeling of brain anatomy. Subdivision of the gyri and exploring differing functional connectivities between these subdivisions with fMRI or SEEG may also help explain why differing seizure semiologies arise from the same gyrus in patients with epilepsy (Mailo and Tang-Wai, 2015).

In contrast to atlases guided by anatomical features, parcellations that are based solely on resting fMRI or diffusion imaging rely on clustering methods that can introduce bias in the shape and size of the clusters. Spectral clustering, for example, is known to be biased towards producing similarly sized clusters (Rokach and Maimon, 2005) while functional subregions are known to vary in size. For example, in the precentral gyrus and calcarine cortex, the subregions supporting hand function and central vision, respectively, are much larger than those supporting leg function or peripheral vision. As such, atlases parcellating the brain into subregions should account for this variation.

Last, it should be noted that most popular software packages (e.g. FreeSurfer (Fischl, 2012), BrainSuite (Shattuck and Leahy, 2002), BrainVISA (Cointepas et al., 2001)), use coregistration based purely on structural (T1/T2 weighted) images. Therefore, it is reasonable to expect stronger consistency across subjects after registration when anatomical landmarks are also used in the construction of the atlas itself.

In this paper, we present both anatomical (BCI-DNI atlas) and hybrid anato-functional (USCBrain atlas) atlases intended for use in segmenting brain MRI data. The BCI-DNI atlas was prepared using a high-resolution 3D T1-weighted (T1W) MRI images of a single subject. The atlas was meticulously extracted and anatomically manually-labeled by a neuroanatomist based on known sulcal and gyral landmarks (Damasio, 2005; Pantazis et al., 2010). It was designed to be used with robust registration algorithms based on volume data (FSL, SPM), cortical surface (FreeSurfer, BrainVISA), or combined surface-volume alignment (e.g. BrainSuite or the CVS program in FreeSurfer).

Additionally, we have developed a principled approach to sub-parcellating anatomically-defined cortical ROIs based on resting fMRI (rfMRI) data. The intended goal of the resulting USCBrain atlas is to define functionally distinct subdivisions of the gyri defined in the BCI-DNI atlas. In cases where resting fMRI data are available for each subject, individualized sub-parcellations can be identified in the absence of anatomical landmarks. However, in many studies, such data are not available, in which case sub-parcellation of gyri can be based on individual anatomical images in combination with group studies of the relationship between rfMRI-based connectivity and cortical anatomy as described here.

We used resting fMRI data to compute connectivity between neighboring vertices to identify boundaries between functionally distinct regions within each gyral ROI. At very fine scales, we expect the boundaries of these functional regions to vary significantly with respect to individual sulcal anatomy (Miller et al., 2009; Ono et al., 1990). We, therefore, limit the number of subdivisions of each gyrus (≤ 4) based on a Silhouette score (Rousseeuw, 1987) and use an independent data set from 60 subjects to evaluate the consistency of the resulting subdivisions across subjects. Each label was named by combining their anatomically defined ROI and their location within the ROI (e.g. anterior-, middle- and posterior-cingulate gyrus). The fMRI analysis and parcellation were performed on the cortical surface. The cortical surface parcellation was then mapped back to the original volume resulting in an atlas with mutually consistent volumetric and cortical surface labels. A probabilistic atlas was also generated for the USCBrain atlas. In addition to the anatomical and functional parcellations, we also provide a delineation between sulcal valleys and gyral crowns which offer an additional set of 26 sulcal subregions per hemisphere. Both parcellations in the atlas have general applicability to cognitive neuroscience and clinical research studies. The BCI-DNI and USCBrain atlases are included in the standard distribution of our open-source software, BrainSuite, and available for download for use with FreeSurfer and FSL (http://brainsuite.org/atlases).

## 2. Materials and Methods

### 2.1. The BCI-DNI Anatomical Brain Atlas

#### 2.1.1. Image Acquisition

A high-resolution 3D MPRAGE image (TE=4.33 ms; TR=2070 ms; TI=1100ms; Flip angle=12 degrees; resolution=0.547mm **×** 0.547mm **×** 0.800mm) was acquired on a 3T Siemens MAGNETOM Trio using a 32-channel head coil. Fat suppression was achieved using spectrally-selective excitation of the water protons. Data was acquired 5 times and averaged to improve SNR at this resolution. The subject is a typical right-handed woman in her mid-thirties. Her brain is midway on the spectrum between dolichocephalic and brachycephalic with no obvious anomalies.

#### 2.1.2. Brain Extraction

T1-weighted images were processed using BrainSuite in a semi-automated fashion to classify tissue types, and to extract and render the surfaces of the inner, mid and pial cortices. Nonuniformity was corrected by both automatic processing and manual guidance to maximize grey-white tissue contrast and ensure accurate tissue classification. Manual corrections were performed on (i) the cortical boundaries to remove inclusion of meninges or exclusion of cortex, (ii) the occipito-cerebellar boundaries to avoid inclusion of cerebellum or tentorium or exclusion of the occipital tissue, and (iii) the grey-white boundaries to preserve fine sulcal and gyral detail.

BrainSuite allows for the processing of imaging data in native resolution which allows for the retention of fine anatomical details for brain extraction and coregistration to our new high-resolution atlas. The software produces tessellated cortical surfaces in the same coordinates as voxelated volumetric labels giving a one-to-one correspondence between brain surfaces and volumes. For registration, this means that surface curvature can guide volumetric alignment of cerebral cortex (Joshi et al., 2007). For analysis, this ensures consistency between surface-based and volume-based measurements. For example, ROI-wise cortical thickness and grey matter volume would be measured within the same boundaries. The cortical surface mesh produced by BrainSuite contains ~350k vertices for the current atlas compared to ~150k vertices rendered for images sampled at 1mm^3^.

#### 2.1.3. Anatomical Labeling

Anatomical labeling was performed manually on coronal single-slice images. Sixty-six cortical regions of interest (ROIs) were delineated manually by H.D. using sulcal and gyral landmarks for guidance as detailed in (Damasio, 2005), a human brain atlas text that is a guide to the localization of brain structure and illustrates a wide range of neuroanatomical variations. H.D. has several decades of experience in the field of neuroanatomy and functional localization in the brain by lesion detection and fMRI studies. Volumetrically, deep gyral boundaries were defined on the atlas based on the two opposing sulcal edges of the gyrus from the coronal view. Cortical volume labels were then interpolated and transferred to the mid-cortical surface meshes for the left and right hemispheres. Cortical surface labels were refined so that label boundaries would follow the sulcal fundi more closely using the geodesic curvature flow method described in (Joshi et al., 2012a) and were then transferred back to the volume label set for consistency between surface mesh labels and volumetric labels. A final manual label editing of the labels was performed on both the cortical surface and volumetric labels. The 29 non-cortical regions including subcortical nuclei, hippocampus, amygdala, corpus callosum, ventricles, brainstem, cerebellum were all labeled manually with no automated refinements. Figure 1 shows the color-coded BCI-DNI atlas with labels and Table A1 in the Appendix lists the full set of ROIs labeled in the atlas.

**Figure 1.**
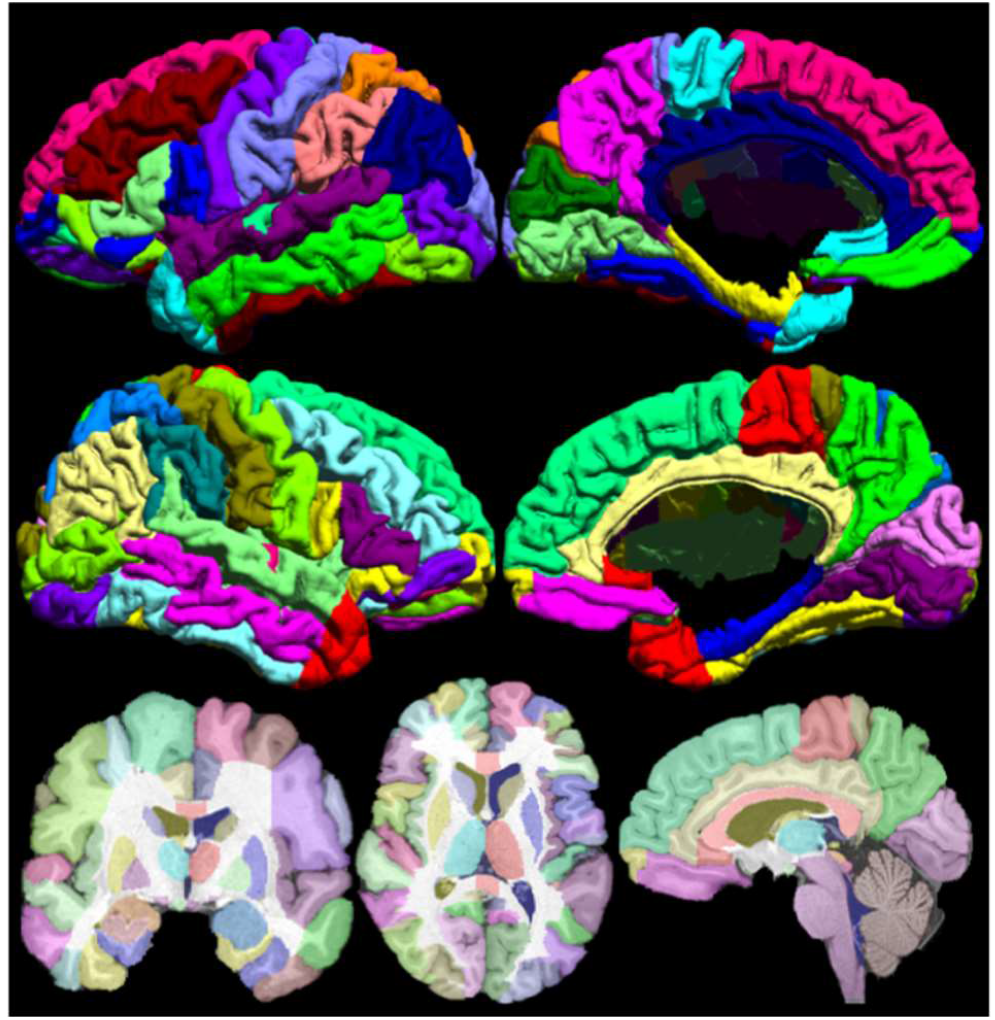
The BCI-DNI anatomical atlas with 95 regions of interest (66 cortical, 29 noncortical) manually labeled by an expert neuroanatomist. Labeled left (top row) and right hemisphere (middle row) of the lateral (left column) and mesial (right column) mid-cortical surfaces (bottom row). Single-slice skull-stripped MPRAGE image with labels overlaid on coronal (left), axial (middle) and sagittal (right) orientation.

Twenty-six sulcal curves were delineated on the midcortical surfaces using a sulcal tracing protocol (http://neuroimage.usc.edu/CurveProtocol.html) (Damasio, 2005; Duvernoy, 1999; Pantazis et al., 2010). Additional sulci were marked for a total of 39 sulci on the left hemisphere and 37 on the right hemisphere (Table A2 in the Appendix). While the 26 sulcal curves are most consistently found on all normal subjects on both cortical hemispheres, the 13 (right) and 11 (left) additional sulci are not included in our sulcal tracing protocol but are still commonly found.

We also delineated sulcal regions around the 26 sulcal curves for applications where users want to differentiate between sulcal valleys and gyral crowns. The delineations of the sulcal regions were performed for each of the atlas’s cortical hemispheres on the mid-cortical surface meshes. The mid-cortical surface representations were smoothed and mean curvature was computed. For this purpose, principal curvatures were computed by fitting quadratic polynomials in each of the vertex neighborhoods (Desbrun and Polthier, 2007). The mean of the two principal curvatures was then computed (Do Carmo, 2016). Sulcal regions were defined as those with negative mean curvature found by thresholding on the zero-level set. The 26 sulcal traces were then transferred to the surfaces. Any interruptions in the sulcal regions along each of the sulci were corrected using nearest neighbor labeling relative to the sulcal curve. Finally, the 26 sulcal regions were identified using connected component analysis for each cortical hemisphere. The process is illustrated in Figure 2.

**Figure 2:**
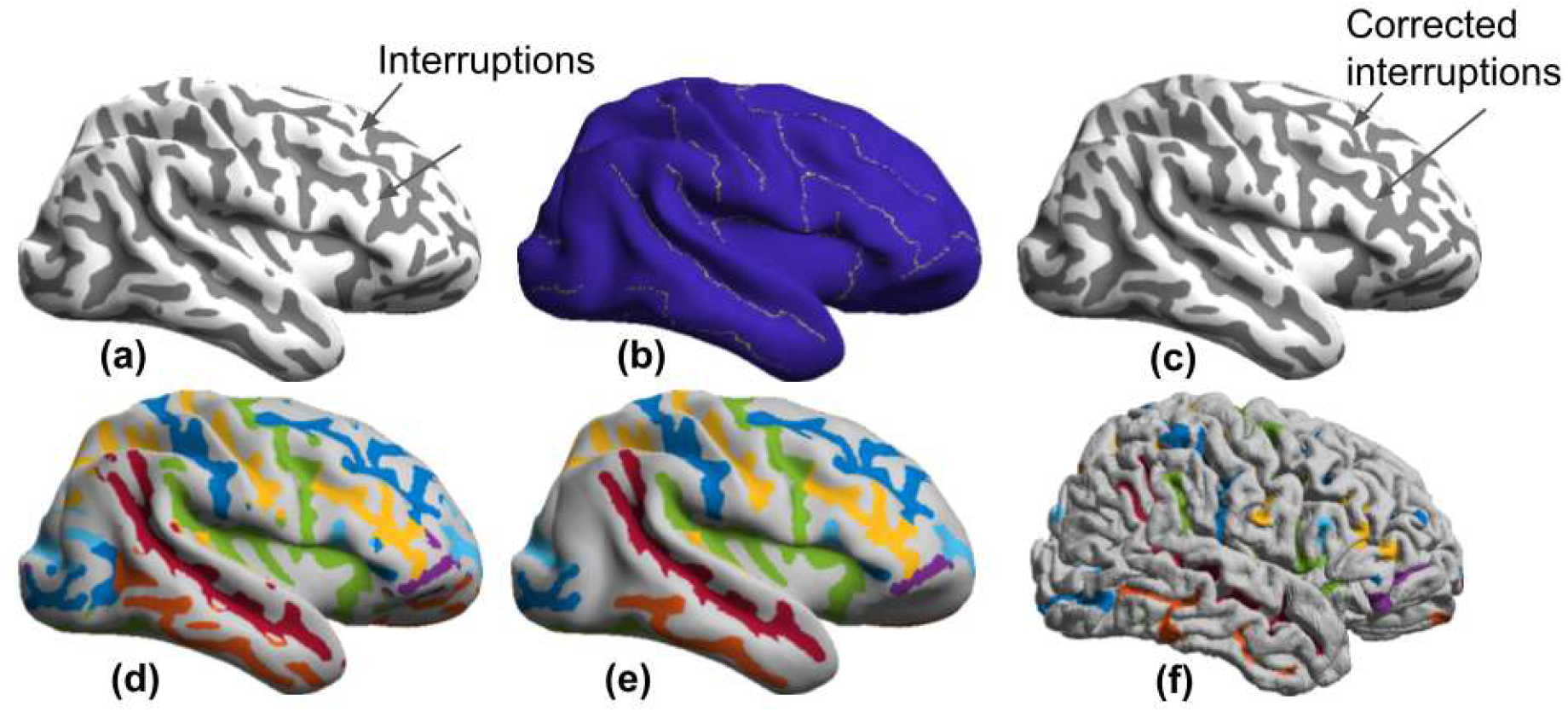
The procedure for delineation of sulcal regions: (a) first, mean curvature maps were computed and thresholded. (b) The 26 sulcal curves were transferred to the surface. (c) These were used to correct for the interruptions along the sulcus in the curvature maps. (d) The 26 sulcal curves were used to label the individual sulcal regions. (e) These regions were further refined using connected component analysis and morphological smoothing. (f) The same sulcal regions as in (e) shown on the original mid-cortical surface.

The final BCI-DNI atlas consists of a set of 95 regions of interest (66 cortical, 29 noncortical) defined with respect to the volumetric space. Also included is a surface-based atlas in which the same 66 gyral (cortical) regions are delineated and labeled in addition to identification of 76 sulcal curves and surface delineation of sulcal banks.

### 2.2. The USCBrain Hybrid Brain Atlas

The anatomical parcellation defined by the BCI-DNI atlas was subdivided using rfMRI data from the Human Connectome Project (HCP). For this purpose, the BCI-DNI atlas labels were transferred to the HCP grayordinate space (Glasser et al., 2013) in which the HCP’s minimally preprocessed rfMRI data are available. After sub-parcellation, the labels were transferred back to the original BCI-DNI atlas space.

#### 2.2.1. Study Population and Data Preparation

A 40 subject dataset (healthy adults, ages 22-35, 16 males, 24 females) was obtained from the WU-Minn Human Connectome Project (HCP) database (Glasser et al., 2013; Van Essen et al., 2013). T1W images and four sessions of rfMRI data were utilized for each subject (TR=720ms, TE=33.1ms, 2mm isotropic voxels, scan time= 15mins). All data was processed with the HCP’s minimal pre-processing pipeline (MPP) (Glasser et al., 2013). The pre-processing pipeline includes processing of T1W images of each subject using FreeSurfer (Fischl, 2012) for the identification of cortical surfaces and coregistration of the surfaces to a common atlas. These surfaces were then down-sampled to a common standard cortical surface mesh (32K Conte-69). Resting fMRI data were corrected for acquisition artifacts and subject to a non-aggressive spatiotemporal filtering as described by (Glasser et al., 2013). The time-series data were then linearly resampled onto the mid cortical surfaces generated by FreeSurfer and transferred to the 32K Conte-69 grayordinate representation (Glasser et al., 2013). We additionally denoised the rfMRI data by applying nonlinear temporal nonlocal-means (tNLM) filtering (Bhushan et al., 2016). The tNLM filtering reduces local signal fluctuations in the fMRI data without the spatial blurring that occurs with standard linear filtering. The resulting time series at each vertex were normalized to zero mean and unit norm.

The BCI-DNI atlas was processed with the FreeSurfer pipeline to generate cortical meshes for inner and pial surfaces and to coregister these meshes to the fsaverage brain atlas in FreeSurfer using spherical mapping and curvature-based registration. The BCI-DNI labels from the inner BrainSuite cortical surface were transferred to the inner FreeSurfer cortical surface mesh. This transfer is possible because both software packages generate very similar cortical surfaces. For both cortical hemispheres, using the coregistered spherical maps provided with the HCP data, the labels from the BCI-DNI atlas were transferred to the 32K Conte-69 surface meshes by nearest neighbor interpolation. These spherical maps are used as intermediate representation for cortical surface representation by FreeSurfer as part of the HCP pipeline (Fischl et al., 1999). As a result, for each subject a BCI-DNI-atlas labeled 32K Conte-69 mesh in the grayordinate coordinates was obtained, the same space in which the HCP rfMRI data is also available.

#### 2.2.2. Resting fMRI Based Sub-Parcellation

For each ROI from the BCI-DNI atlas in the grayordinate representation (Figure 3(a, b)), we computed a similarity matrix for each subject using the rfMRI data. The similarity measure was computed between each pair of vertices in the ROI using their respective time series (*X, Y*) as *s*(*X, Y*) = *π* − cos^−1^(*X* · *Y*), where the dot-product *X* · *Y* indicates the Pearson correlation coefficient between the time-series and cos^−1^(·) represents the principal value of the inverse cosine function. We chose this measure because cos^−1^(*X* · *Y*) in the range [0, *π*) represents the geodesic distance between the two unit-length vectors *X* and *Y* and is therefore a true metric on the hypersphere. This pairwise similarity matrix was input to the spectral-clustering normalized graph-cut algorithm (Shi and Malik, 2005, Craddock et al., 2013) to subdivide each ROI. The spectral clustering algorithm chooses a set of graph cuts defining the region boundaries that minimize the ratio of the total edge weight along with the cuts to the total edge weight within regions. As a result, the gyrus is subdivided such that the subdivisions have distinct connectivity profiles.

**Figure 3.**
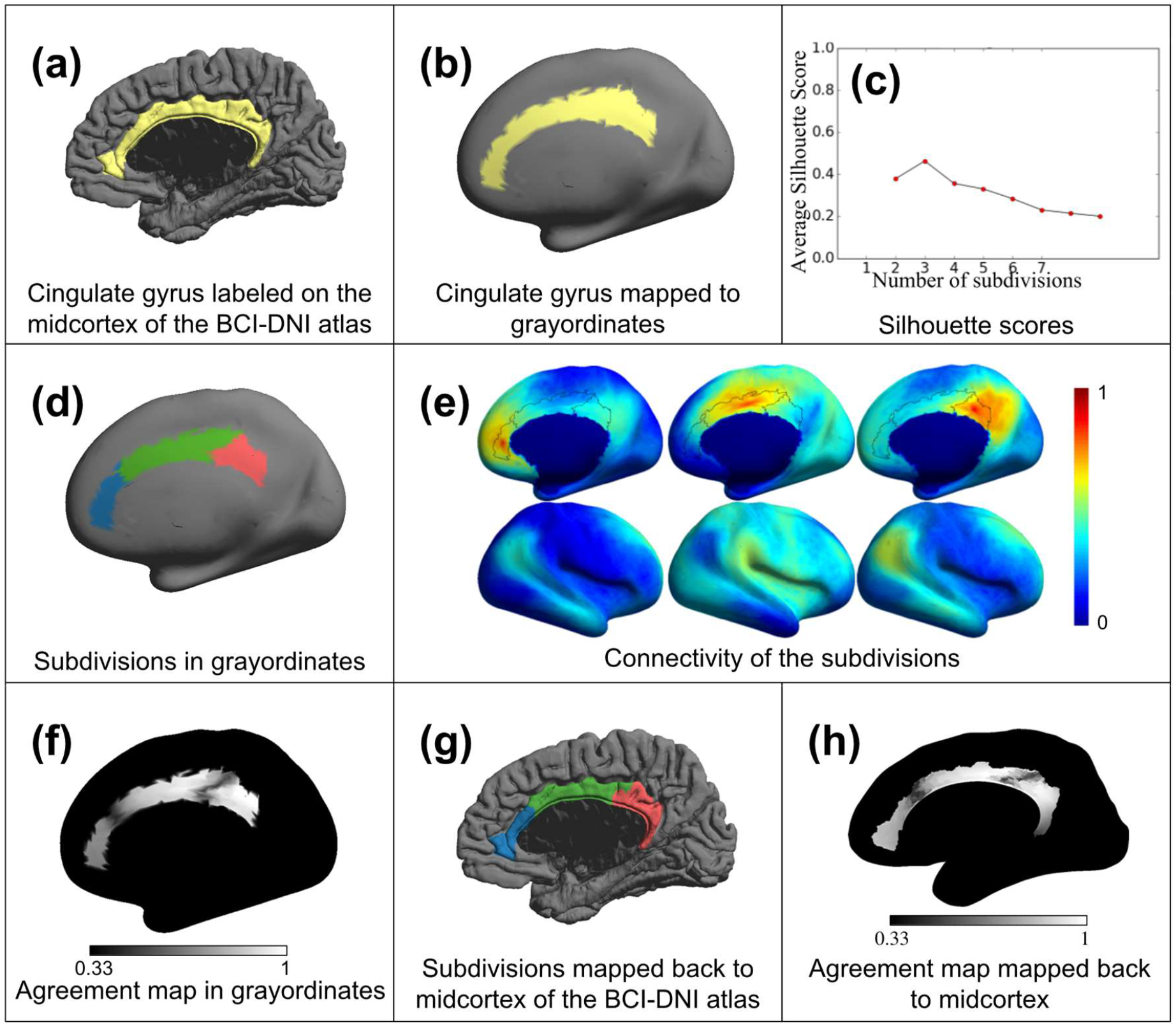
The process of subdivision is shown in the figure with right cingulate as an illustrative example: (a) right cingulate gyrus from BCI-DNI atlas; (b) label transferred to the grayordinate space of HCP in which the fMRI data is available; (c) Silhouette score computed for different numbers of clusters; (d) subdivisions performed for one of the 40 subjects; (e) seed-based connectivity using centroid vertex of each subdivision is computed to show differences in connectivity of each subdivision; (f) agreement maps across 40 subjects; (g),(h) the subdivision labels and agreement maps transferred back to the midcortex of the BCI-DNI brain.

To determine the optimal number of subdivisions, we computed the Silhouette score (Rousseeuw, 1987). This measure compares the similarity of each vertex to other vertices in its own cluster in comparison to all other clusters and can be briefly described as the normalised difference between mean inter- and intra-cluster distances. It has been previously proposed as a metric to automatically choose the number of different clusters in general clustering applications (de Amorim and Hennig, 2015). Silhouette scores vary from −1 to +1, where a large positive value indicates that the vertex matches the assigned cluster well and a small or negative value indicates a poor fit. The average Silhouette score over all subjects for a different number of subdivisions N was computed and the N that maximized the average Silhouette score was chosen as the number of subdivisions for each anatomical ROI. For example, Figure 3(c) indicates an optimal value of N=3 for right cingulate.

The sub-parcellations were computed for 40 HCP subjects. The cluster labels assigned to gyral subdivisions by the clustering algorithm have an arbitrary order and are not consistent across subjects. We used the Hungarian algorithm (Kuhn, 1955) to reorder the cluster labels so that they were maximally consistent across subjects. The final subparcellation of the ROIs was generated by taking a majority vote (random assignment in case of ties) of labels at each vertex over the 40 subjects (Figure 3(d)). Qualitative verification of the subparcellation was performed by visualizing the functional connectivity patterns between subparcellations of each anatomical ROI. The geometric centroid vertex within a cluster was used as the seed point to compute the correlation of its time series to that of all other vertices throughout the brain. Correlations were averaged across subjects and displayed as illustrated in Figure 3(e). To check the consistency of labels across the 40 subjects, we computed a label agreement map as follows. At each vertex, we performed pairwise comparisons of labels between each pair of subjects and assigned a value of 1 if they are same or 0 if they are different. The agreement map was then computed as the average over all possible pairs, consisting of values in the range (*1/N,1*) where *N* is the number of subdivisions. High agreement scores were expected towards the center of the labels while low scores were expected near the edges. Sharp boundaries between subdivisions are indicative of consistency of subdivisions across subjects and were deemed preferable. A grayscale modulated agreement map is shown for right cingulate in Figure 3(f). We then mapped the labels and agreement maps for each of the subdivided gyri from the 32K Conte-69 inner surface mesh back to the BCI-DNI atlas inner cortical surfaces using nearest neighbor interpolation as described in section 2.2.1 (Figure 3(g,h)). The inner, mid cortical and pial surfaces generated by BrainSuite have the same number of vertices that are in correspondence with each other. Therefore, labels were copied directly from the inner cortical surface to the mid and pial surfaces.

We note that the optimal number of regions was identical for homologous ROIs in the left and right hemispheres in most cases. For example, for cingulate gyrus (Figure 3) the Silhouette scores had a maximum of 3 subdivisions for both hemispheres. However, in some ROIs, the optimal number of left and right subdivisions differed, specifically in the angular, middle occipital, middle temporal, and superior temporal gyri. The angular gyrus (Figure 4) showed maximum Silhouette scores for 2 (left) and 3 (right) subdivisions respectively. However, on inspection the connectivity maps were distinct, and the agreement maps also showed distinct boundaries for 3 subdivisions in both hemispheres. Therefore, the angular gyrus was subdivided into 3 regions for both hemispheres, as illustrated in Figure 4. A similar process of inspection of the connectivity and agreement maps was used to select equal numbers of subdivisions in left and right homologous regions for the other gyri listed above. In addition, while the inferior temporal gyri had optimal Silhouette scores for 2 subdivisions bilaterally, 3 subdivisions were chosen because of the large size of the gyrus and the fact that connectivity and agreement maps indicated a clear and distinct delineation of three regions across subjects.

**Figure 4:**
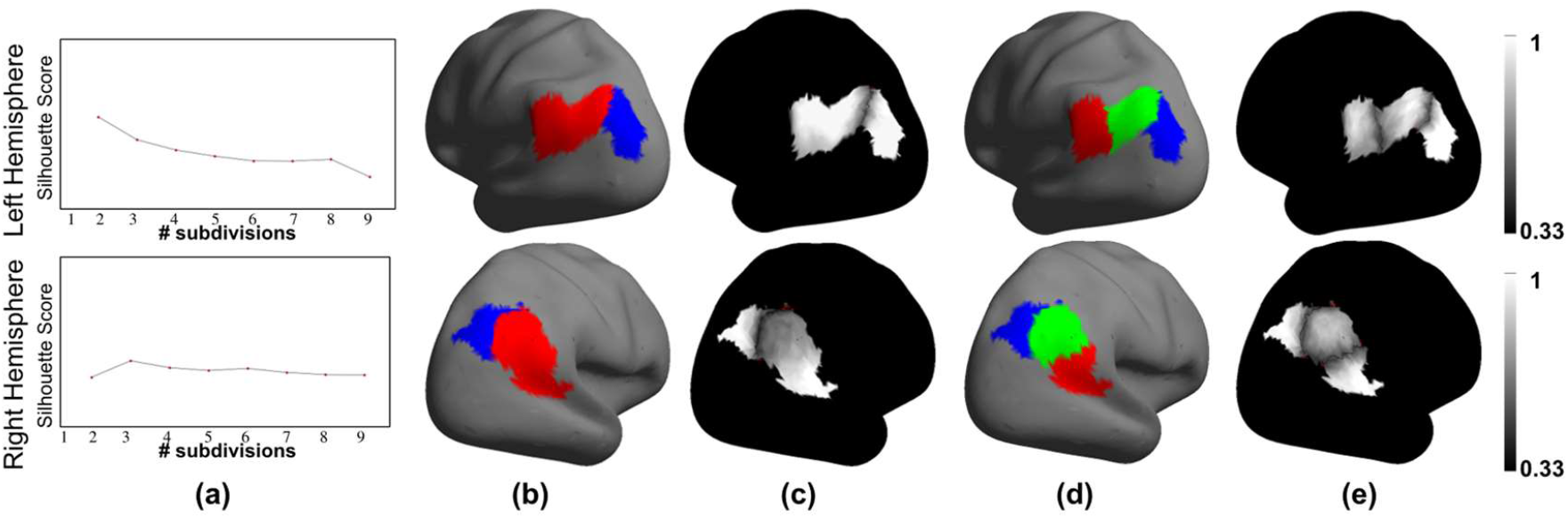
Subdivision of Angular gyrus for both left and right hemispheres: (a) Silhouette analysis; (b) 2 subdivision labels; (c) the corresponding agreement map; (d) 3 subdivision label for the same gyrus and (e) corresponding agreement map; As can be seen, 3 subdivisions resulted in crisper agreement maps for the right hemisphere. For the left hemisphere, while 2 subdivisions showed better agreement maps, 3 subdivisions showed a still acceptable agreement map and high Silhouette score. we therefore choose 3 subdivisions to maintain symmetry between the two hemispheres.

As a final step, the refined cortical surface labels in the BCI-DNI atlas space were propagated from the surface onto the volumetric atlas. Two intermediate surfaces (one each between inner/mid and mid/pial) were generated and labeled giving us a total of 5 labeled surfaces in the cortical ribbon. The gyral volume labels (grey matter, along with the white matter regions) as defined in the BCI-DNI atlas, were filled in by nearest-neighbor interpolation with these surface labels as a reference. The agreement map was propagated to the volume in a similar manner to retain a one-to-one correspondence between the surface and volume labels as shown in Figure 5. Agreement maps were combined across all cortical gyri to generate probability maps that represent inter-subject agreement of the labels across the 40 subjects at each vertex in the cortex, Figure 6.

**Figure 5.**
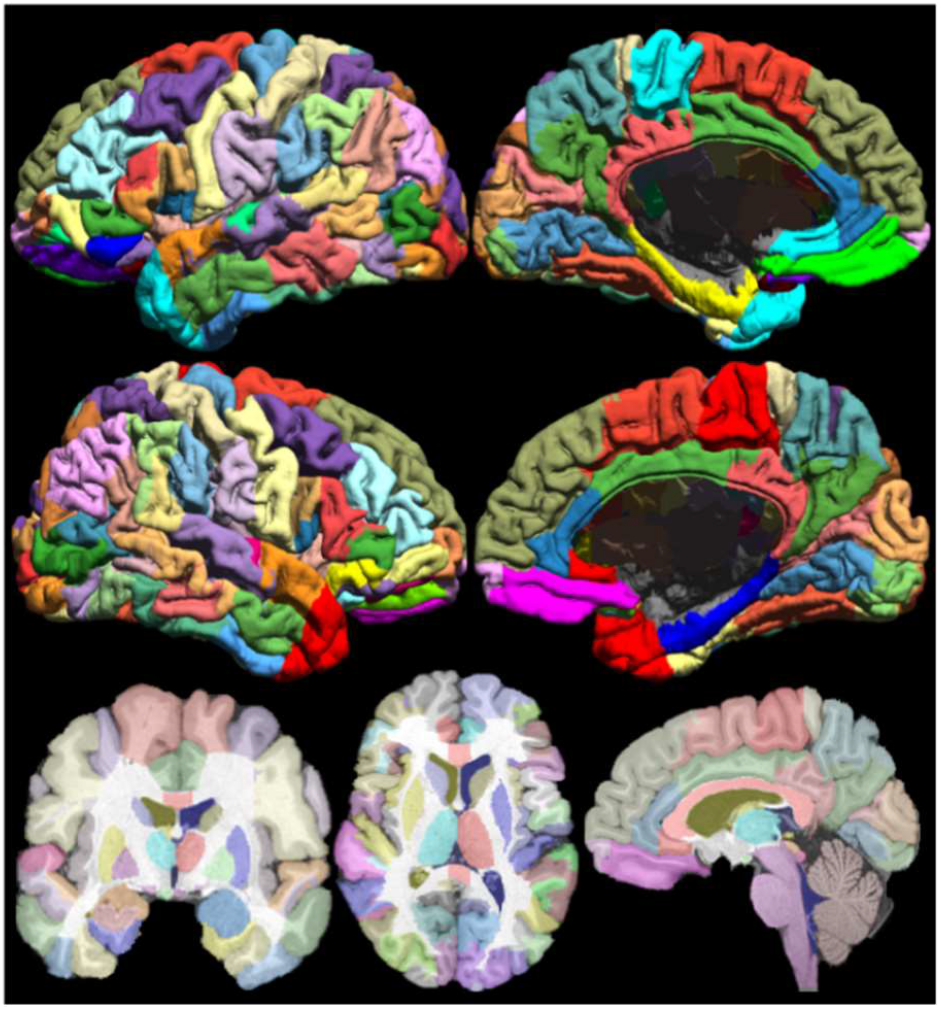
The USCBrain atlas with 130 cortical regions of interests subdivided using rfMRI data. The figure shows the mid-cortical surface of the atlas, color coded for each sub-parcellated region. Labeled left (top row) and right (bottom row) hemisphere and mesial (right column) and lateral (left column) mid-cortical surfaces. (bottom row) Single-slice skull-stripped MPRAGE image with labels overlaid on coronal (left), axial (middle) and sagittal (right) orientation.

**Figure 6.**
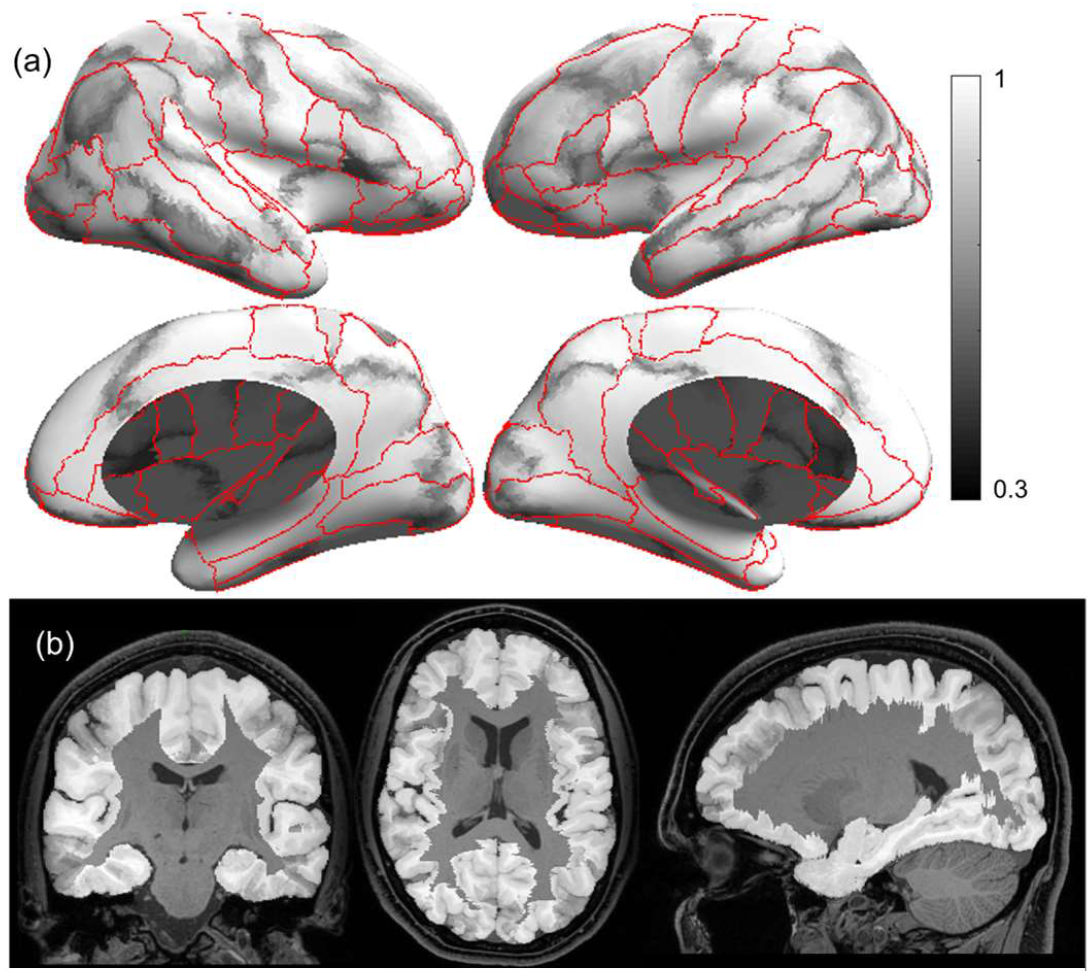
Grayscale modulated plot showing agreement of the USCBrain atlas labels across the 40 subjects. White (value of 1) indicates perfect agreement at that vertex. The probability maps were generated on the surface as shown in (a) and transferred to the volume as shown in (b). Red curves indicate ROI boundaries of the BCI-DNI atlas.

As with the BCI-DNI atlas, the final USC Brain atlas consists of consistent volumetric and surface-based parcellations and labels. The volumetric component of the last atlas contains a total of 159 regions of interest (130 cortical, 29 noncortical) defined with respect to the volumetric space. The surface-based component of the atlas has the same 130 sub-gyral (cortical) regions delineated and labeled in addition to identification of 76 sulcal curves and delineation of sulcal banks for each sulcus.

### 2.3. Validation

In order to evaluate the consistency of labeling of gyral subdivisions using the USCBrain atlas, we investigated how precisely the boundaries of the sub-parcellations could be identified by coregistration of the atlas to subject using T1W MRI data only. The rationale for using the USCBrain atlas-based parcellation instead of individual rfMRI based parcellation is that in most imaging studies, T1W images are acquired with enough anatomical detail for robust registration. Conversely, resting fMRI data are not routinely collected, and even when available are rarely of the quality of the HCP data which is required for reliable individual functional parcellation. We, therefore, investigate whether the cortex can be sub-parcellated into functionally meaningful regions using anatomically driven coregistration alone.

An additional separate set of 60 HCP subjects, with T1W images and four 15-minute rfMRI sessions, were selected and processed as described above. We compared consistency between (a) sub-parcellations by atlas coregistration to the individual T1W image as described in section 2.2.1 and (b) sub-parcellations for that same individual obtained using each of the four sessions of rfMRI data for each subject as described in section 2.2.2. We then compared the sub-parcellation results from (a) to the results from (b) using the Adjusted Rand Index (ARI) (Rand, 1971) for each of the 4 sessions and all subjects (60*4=240) for all ROIs. The ARI ranges from 0 to 1 where 0 indicates labeling performance equivalent to random assignment while 1 indicates perfect agreement. We do expect to see intra-subject variability in the functional parcellation for different rfMRI sessions since repeated rfMRI scans within the same subject can capture different connectivity patterns (Miller et al., 2009). The measures of uncertainty in parcellations determined from individual rfMRI data then provide a baseline score with which to compare parcellation based on the registration of the USCBrain atlas. If the ARI between atlas coregistration and rfMRI-based sub-parcellations is comparable to the ARI between different sessions of rfMRI, then registration of the atlas would serve as an appropriate surrogate for individual rfMRI based parcellation.

### 2.4. Interactivity with BrainSuite, FreeSurfer, and FSL

The atlases were created to be used with the BrainSuite software (http://brainsuite.org), FreeSurfer (https://surfer.nmr.mgh.harvard.edu), and FSL (https://fsl.fmrib.ox.ac.uk/fsl/fslwiki). BrainSuite and FreeSurfer both take the subject’s T1w images as input and generate cortical surface representations. These surfaces are mapped to a flat (for BrainSuite) or spherical (for FreeSurfer) space, and coregistered in that space to atlas surfaces (Fischl, 2012; Joshi et al., 2012b). FSL on the other hand does not generate cortical surface representations, but performs the whole brain volumetric registration using 3D nonlinear registration (Andersson et al., 2007). BrainSuite also performs volume registration using a cortically constrained approach (Joshi et al., 2007) so that the cortical surface and volume alignment results are mutually consistent.

The approximate processing time required for registration and labeling of a typical image volume (image size: 128×256×256; resolution: 1.33mm × 1mm × 1mm) using a desktop workstation (speed Intel Xeon model E5630, 24GB RAM) were, for each software package: 1.5 hours for BrainSuite to perform surface and volume registration and labeling; 20 hours for FreeSurfer to perform surface labeling only, and 1.2 hours for FSL (FLIRT+FNIRT) to perform volume labeling. Scripts and instructions for using the atlases with each software package are available at http://brainsuite.org/using-atlases.

## 3. Results

### 3.1. Parcellation

The BCI-DNI anatomical atlas is shown in Figure 1 with the cortical labels on the mid-cortical surface and the volume labels on the sagittal, coronal and axial slices. A full description of the atlas and downloadable files can be found online (http://brainsuite.org/bcidnibrainatlas/). This atlas has 95 ROIs delineated on the volume of which 66 of these regions are cortical subdivisions (33 per hemisphere). The 76 sulcal curves are also included with the atlas.

The USCBrain atlas is displayed in Figure 5. Of the 33 initial anatomical cortical regions per hemisphere in the original BCI-DNI atlas (Table A1 in the Appendix), 23 ROIs were subdivided resulting in a total of 65 regions per hemisphere. The remaining 8 original ROIs did not exhibit sufficient internal variation in functional connectivity in their rfMRI data to justify subdivision. Most anatomical ROIs were subparcellated into 2 or 3 regions. As expected, large anatomical ROIs such as the superior, middle, and inferior temporal gyri, middle frontal gyrus, and cingulate cortex were subdivided into two or more subdivisions while the smaller ROIs such as the temporal pole, paracentral lobule, and Heschl’s gyrus were not subdivided.

### 3.2. Intersubject Label Consistency

A grayscale modulated image showing agreement in labels across the 40 subjects of the sub-parcellated atlas generated by majority vote is shown in Figure 6. The grayscale ranges from black indicating no consistency to white indicating perfect consistency across subjects. These maps show near-perfect consistencies (~1) near the centers of the sub-parcels as well as at the boundaries of anatomical parcels and reduced consistencies (~.5) at the boundaries of the functional sub-parcels. A few regions showed higher variability including the right inferior temporal and angular gyrus along with the bilateral inferior occipital gyri, where values reached as low as 0.7 in the center of the region, indicating substantial functional variability with respect to the sulcal and gyral anatomy that guides coregistration to the original BCI-DNI atlas.

### 3.3. Anatomical vs Functional Labeling

3.4. As noted earlier, we chose an additional set of 60 HCP subjects disjoint from the set of 40 subjects used to generate the atlas. Box plots of the ARIs for each of the cortical ROIs in the BCI-DNI atlas are shown in Figure 7. For each ROI, we show consistency between atlas and rfMRI based subparcellation on the left and consistency between different rfMRI sessions on the right. The ROIs that were not subdivided show perfect agreement with an ARI of 1 but are included in the figure for completeness.

**Figure 7:**
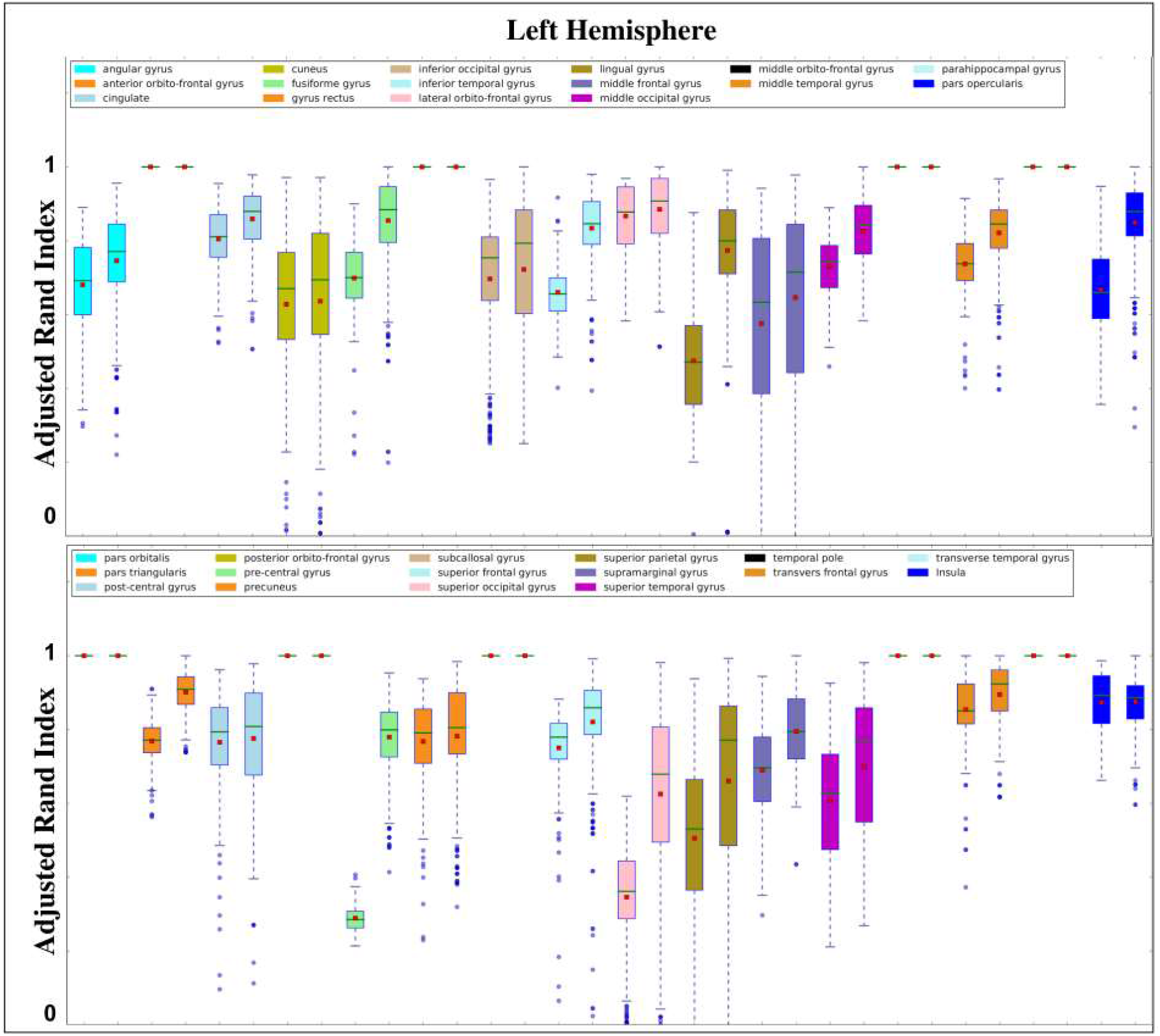
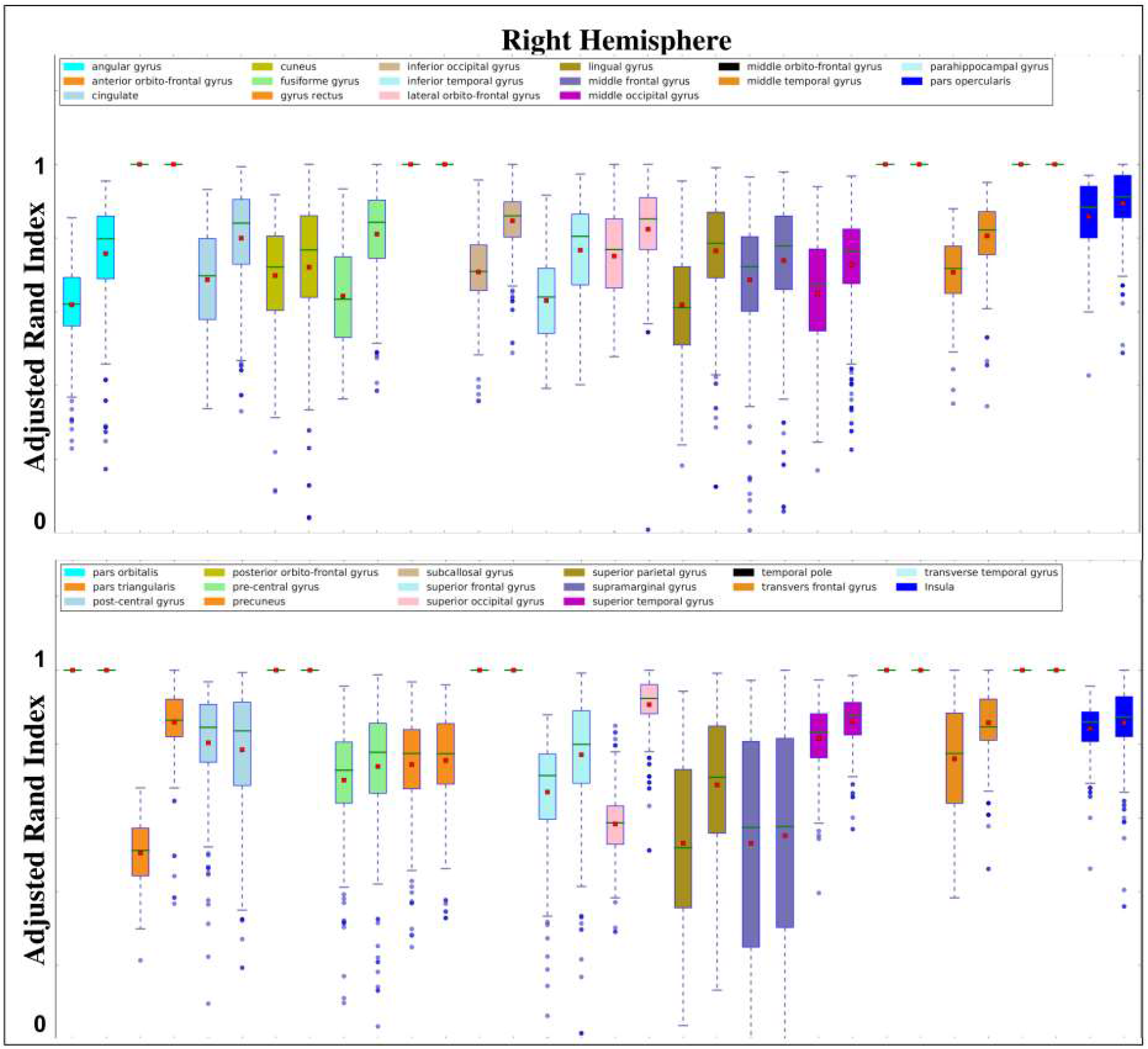
Adjusted rand indices (ARIs) for 60 subjects for left and right hemispheres. Each ROI has two box plots. The first (left) box plot compares atlas based sub-parcellation to rfMRI sub-parcellation. The second (right) box plot compares rfMRI based sub-parcellations across sessions

These figures show that, for the 60 validation subjects, overall the sub-parcellations generated by atlas-based registration relative to rfMRI results show consistently lower average ARI than those based on comparison of individual rfMRIs from different sessions. With the high quality and 15-minute duration of the HCP data, not surprisingly, using an individual subject’s rfMRI data will typically generate a better functional parcellation of that subject than using the atlas However, as noted above, most studies do not routinely acquire comparably high-quality rfMRI data. In this case, the atlas-based subparcellations serve as a surrogate for individual frMRI based parcellation. Figure 7 shows that the majority of regions in the atlas have mean ARIs for atlas vs rfMRI within the inter-quartile range of the rfMRI vs rfMRI results. These results are summarized in Figure 8. For each of the 60 test subjects we computed the ARIs between sub-parcellations of each cortical ROI in the BCI-DNI atlas based on registration to the USCBrain and sub-parcellations computed directly using individual rfMRI data. These were then averaged over the four sessions for all subjects. Figure 8(a) shows the resulting average ARIs for each region. Figure 8(b) shows the equivalent figure for the ARIs between different rfMRI sessions averaged over all possible session pairs for each subject and overall subjects. Finally, in Figure 8(c) we show Cohen’s D statistic comparing means normalized by the pooled standard deviation. These images indicate the consistency of our parcellation is particularly poor in left and right pars triangularis and right middle-temporal gyrus with Cohen’s D >1. Left lingual gyrus, left precentral gyrus, and left fusiform gyrus also show larger values but all <1. Elsewhere consistency is far better (see Discussion for more details) These images, together with the results in Figures 7 and the agreement images in Figure 6, can be used to guide users in determining expected reliability of atlas-based identification of subparcellations by gyrus.

**Figure 8.**
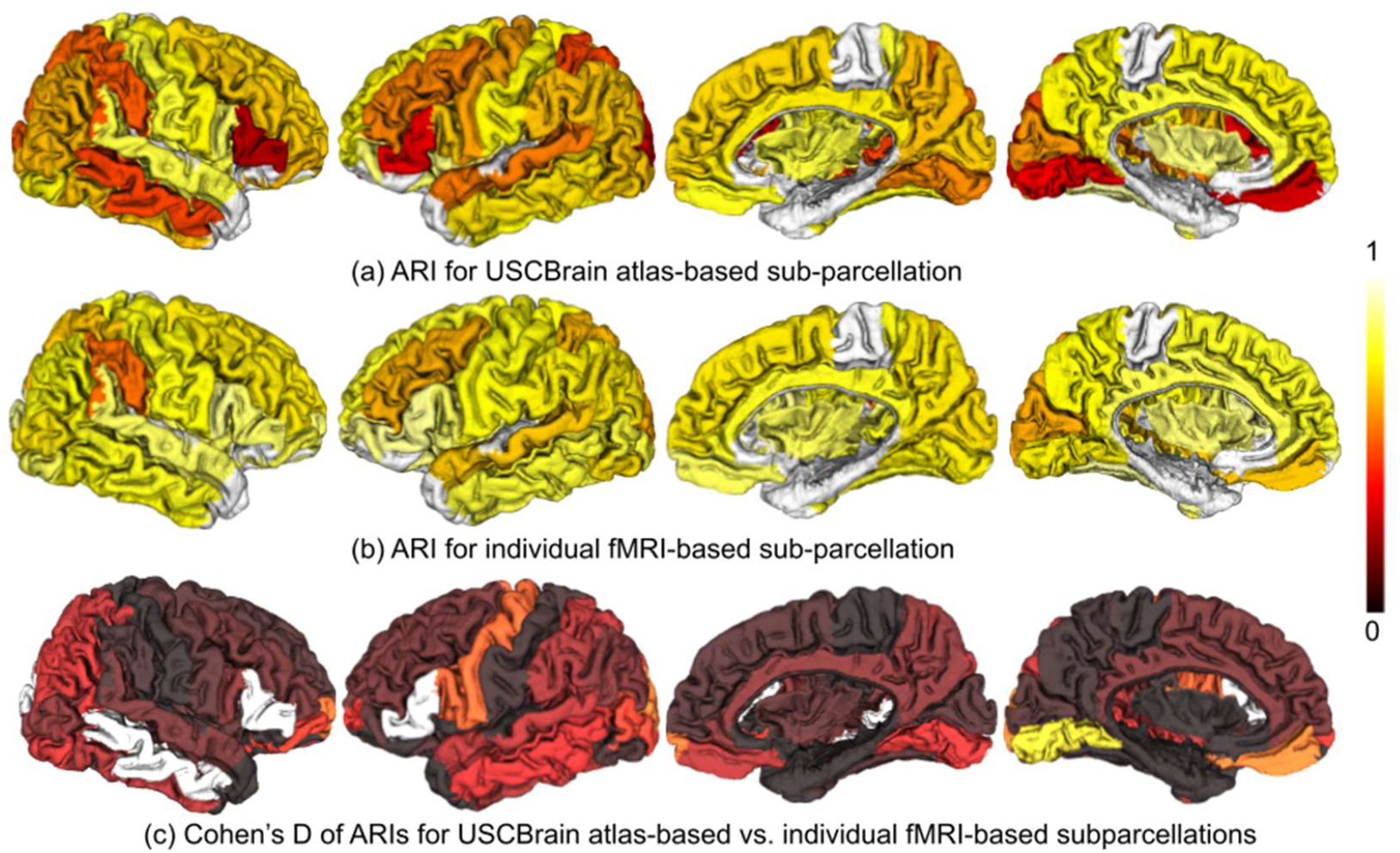
The mean ARI shown as colorcode ROIs between (a) USCBrain atlas based sub-parcellation and fMRI based subparcellation; (b) pairs of fMRI based subparcellation from 4 different scan sessions for each individual. (c) The Cohen’s D statistic computed as the difference in means between (a) and (b) scaled by their pooled standard deviation.

### 3.5. Comparison to other atlases

We compared the properties of the USCBrain and BCI-DNI atlases with a set of nine commonly used cortical parcellations (see Table 1 for details) in terms of functional homogeneity and agreement with cytoarchitectonic boundaries. All atlases were evaluated in HCP fs_LR32k surface space (Glasser et al., 2013; Van Essen et al., 2013). For the parcellations that were released only in volume space (Yeo, Gordon, Gordon2), we either used their publicly available surface representations from a previous study (Arslan et al., 2017), or projected to the surface space using the Brainsuite Functional Pipeline (BFP, https://github.com/ajoshiusc/bfp). For the Yeo atlas (Yeo et al., 2011), we used a relabeled version from Yeo’s lab that splits each network into spatially contiguous parcels. For the parcellation described in (Gordon et al., 2016), we included both its official release, denoted as Gordon, which has approximately 30% of the vertices not assigned to any clusters, and the version used in (Arslan et al., 2017), which labels these missing vertices by iterative dilation (denoted as Gordon2; see Table 1). The atlases used in the comparison are arranged into three groups based on whether they were parcellated using structural data only (‘anatomical’), functional only (‘functional’), or multimodal data (‘hybrid’).

**Table 1.** Listed atlases included in the comparison in Figs. 9 and 10. Atlas type: functional: the atlas is generated based on rsfMRI only; anatomical: the atlas is generated based on anatomical landmarks only; hybrid: the atlas uses both functional and anatomical information

| Name | Type | Resolution | Reference | Source |
| --- | --- | --- | --- | --- |
| **Yeo-51**<br>**Yeo-114** | Functional | 51 (L: 26, R: 25)<br>114 (L: 57, R: 57) | (Yeo et al., 2011) | https://github.com/ThomasYeoLab/CBIG/tree/master/stable_projects/brain_parcellation |
| **Power** | Functional | 130 (L: 65, R: 65) | (Power et al., 2011) | https://biomedica.doc.ic.ac.uk/brain-parcellation-survey/ |
| **Schaefer-100/200/300** | Functional | 100 (L: 50, R: 50)<br>200 (L: 100, R: 100)<br>300 (L: 150, R: 150) | (Schaefer et al., 2018) | https://github.com/ThomasYeoLab/CBIG/tree/master/stable_projects/brain_parcellation |
| **AAL** | Anatomical | 97 (L: 47, R: 50) | (Rolls et al., 2020) | https://github.com/ajoshiusc/bfp/blob/main/supp_data/AALv3_grayordinate_labels.mat |
| **Desikan** | Anatomical | 70 (L: 35, R: 35) | (Desikan et al., 2006) | https://biomedica.doc.ic.ac.uk/brain-parcellation-survey/ |
| **Destrieux** | Anatomical | 150 (L: 75, R: 75) | (Fischl et al., 2004) | https://biomedica.doc.ic.ac.uk/brain-parcellation-survey/ |
| **Gordon**<br>**Gordon2** | Functional | 333 (L: 161, R: 172) | (Gordon et al., 2016) | https://balsa.wustl.edu/WK71<br>https://biomedica.doc.ic.ac.uk/brain-parcellation-survey/ |
| **Brainnetome** | Hybrid | 210 (L: 105, R: 105) | (Fan et al., 2016) | https://biomedica.doc.ic.ac.uk/brain-parcellation-survey/ |
| **Glasser** | Hybrid | 360 (L: 180, R: 180) | (Glasser et al., 2016) | https://balsa.wustl.edu/study/show/RVVG |

#### Functional homogeneity

We used resting fMRI data from 158 subjects in the ADHD-200 dataset (http://fcon_1000.projects.nitrc.org/indi/adhd200/). The rsfMRI data first was preprocessed with Brainsuite’s BFP pipeline (https://github.com/ajoshiusc/bfp), which includes motion correction, skull stripping, grand mean scaling, temporal filtering, detrending, spatial smoothing, nuisance signal regression, and GPDF filtering (Li et al., 2020, 2018). We then computed the group-average functional connectivity between all pairs of vertices on the cortical surface as the Pearson correlation between their respective time series, averaged over all subjects and transformed using Fischer-Z. Next, the functional homogeneity *ρ*_i_ of cluster *i* was computed by averaging the functional connectivity values over all vertex-pairs within each cluster. Finally, to account for different cluster size distributions between parcellation schemes, we followed (Schaefer et al., 2018) and computed the weighted functional homogeneity as a global measure, defined as

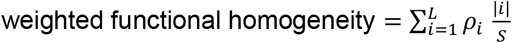

where |*i*| is the number of vertices in cluster *i* and *S* is the total number of cortical vertices. We also computed weighted functional homogeneity variance as 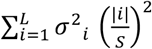, where *σ*^2^_*i*_ is the variance in cluster *i*. The results of this comparison are shown in Fig. 9.

**Figure 9:**
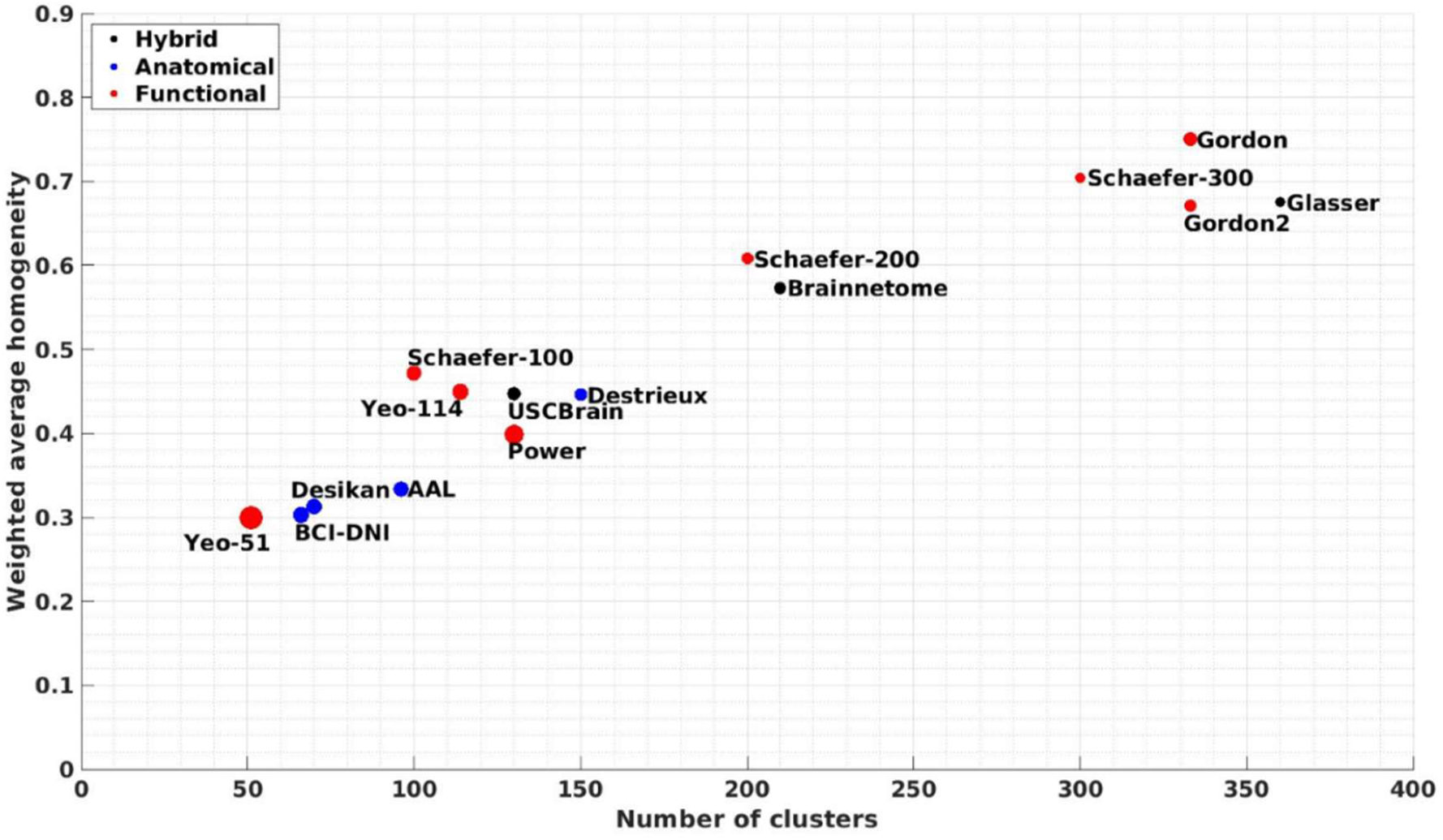
Weighted average functional homogeneity. The diameter of each circle is proportional to weighted functional homogeneity variance.

#### Cytoarchitectonics

To evaluate the agreement of different parcellations with Brodmann cytoarchitectonic areas we used the method and script described by (Arslan et al., 2017) (https://github.com/sarslancs/parcellation-survey-eval) that measures the overlap of a parcellation with selected regions of the Brodmann atlas as provided by the Human Connectome Project (Van Essen et al., 2013). These Brodmann areas consist of the primary somatosensory cortex, primary motor cortex, premotor cortex, Broca’s area, visual cortex, and perirhinal cortex. For each Brodmann area, the script computes Dice similarity coefficients with respect to each parcellated atlas by first matching and merging parcels to find the group that jointly provides best agreement, in terms of the Dice coefficient, between combined parcels and the Brodmann area. A global score is then computed by averaging Dice coefficient scores for these merged parcels for all 8 of the selected Brodmann areas.

This merging process allows comparison of atlases with widely varying numbers and sizes of parcels. The results of this comparison are shown in Fig. 10.

**Figure 10.**
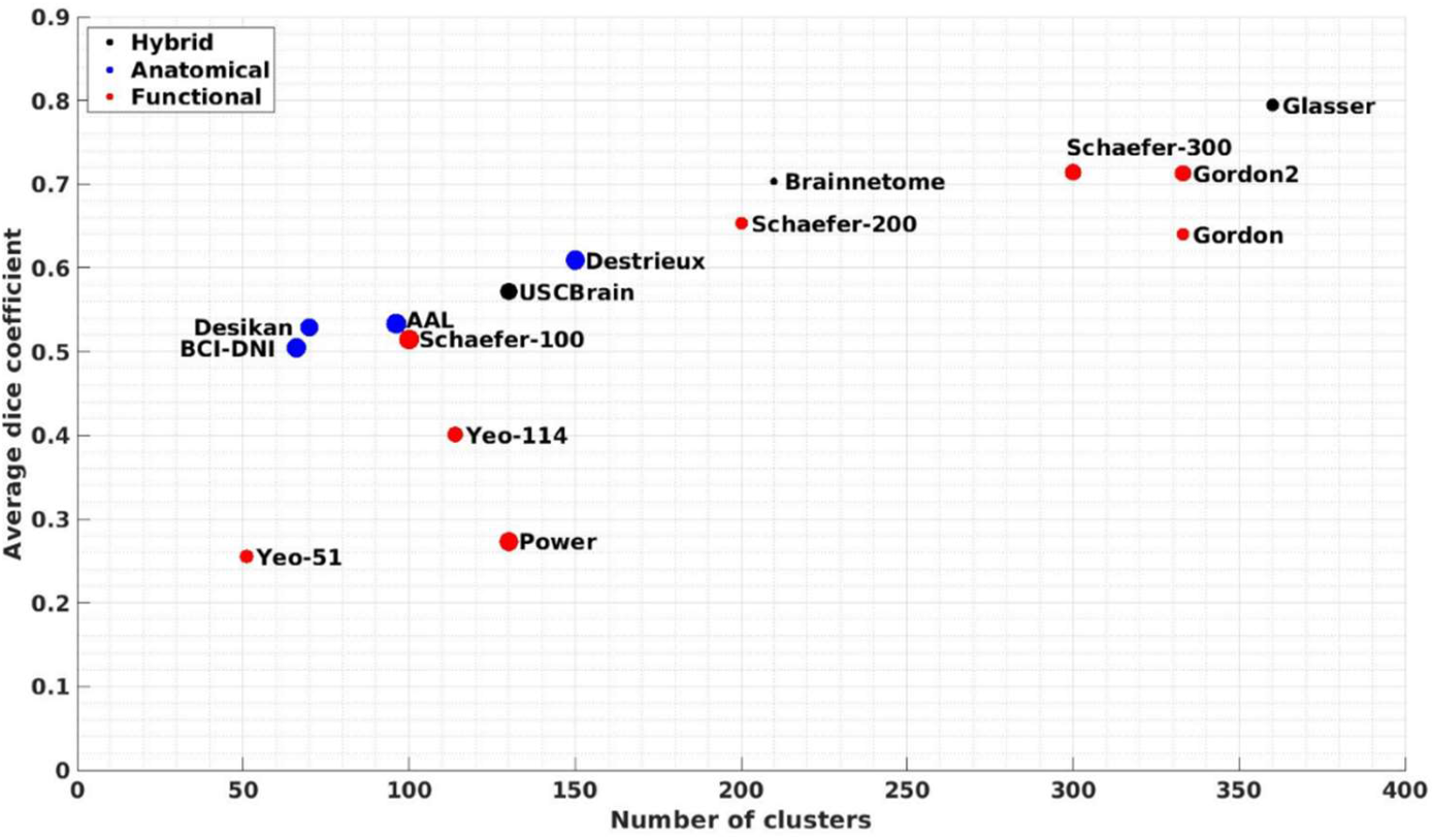
Agreement with cytoarchitectonics. Average Dice coefficients are computed between merged parcels (as described in the text) and each of the eight Brodmann regions. The diameter of each circle is proportional to the variance of dice coefficient across the eight regions.

## 4. Discussion

The aim of this work was to develop a new brain atlas that has two levels of parcellation. The anatomical BCI-DNI atlas provides a parcellation of cortex based on established anatomical sulcal and gyral patterns. The hybrid USCBrain atlas then provides a refined parcellation that attaches subgyral labels based on functional connectivity using resting fMRI data for 40 subjects from the Human Connectome Project (HCP). Both atlases include consistent volumetric and surface-based parcellation and labeling. Particular attention was paid to the image acquisition, processing, and labeling methods to capture fine anatomical details, accommodating for the high-quality data increasingly common in recent imaging studies. Special attention was also paid to the selection of the subject used in this atlas as the individual imaged for the commonly used Colin27 stereotaxic atlas (Holmes et al., 1998) has a unique/unsual left lower precentral region. The precentral gyrus, on the pial surface, seems to terminate early before it reaches the Sylvian fissure. This causes inconsistent continuity of the inferior frontal gyrus and the complex of pre- and post-central gyri. This unique/unusual feature causes systematic misidentification of the inferior frontal gyrus and the inferior regions of the pre- and post-central gyri making it a suboptimal target for a template. In contrast, the individual’s brain in the BCI-DNI atlas shows normal architecture, including sulcal and gyral patterns, without unique/unusual features that may bias individual- and group-level parcellations.

Subdelineations of the gyri in the BCI-DNI atlas to form the USCBrain atlas were determined according to measurements of rfMRI connectivity. Parcellations of individual subjects yielded reasonable results in our validation studies based on 60 additional HCP subjects (Figures 7, 8) which establish the level of consistency of these functional boundaries for each region. In order to quantify expected intersubject variability, we also provide a probabilistic agreement map (Figure 6).

### 4.1. Brain Asymmetry

A degree of asymmetry was observed between the right and left hemispheres in both the curvature patterns that determined the boundaries in the BCI-DNI atlas (Figure 1) as well as the functionally driven boundaries in the USCBrain atlas (Figure 4). Notably, the functional boundaries subdividing the angular gyrus, and its surrounding connected gyrus, as well as the middle and inferior temporal gyrus were more asymmetric than other cortical gyri observed based on the topographical location of the boundaries. Regions of the medial and dorsolateral prefrontal cortex, including the cingulate gyrus and precentral gyrus, were less asymmetric. However, all pairs of homologous regions in the brain showed some degree of asymmetry (homologous boundaries varied from one hemisphere to the other by from 1mm to ~1cm) as would be expected given known differences in left-right functional specializations. Symmetry in the number of clusters each gyrus was subdivided into was enforced in the USCBrain atlas. In all four gyri where the Silhouette analysis gave a differing number of subdivisions for left and right hemispheres, reasonable results in terms of distinct connectivity patterns, good intersubject agreement, and Silhouette values relative to their maximum were observed after matching the number of subdivisions as described in Section 2.2.2.

### 4.2. Parcellation Density and Comparison to Other Atlases

Silhouette scores (section 2.2.2) were the main factor that we used to determine the optimal number of subdivisions for each cortical gyrus. Most large ROIs, such as the cingulate gyrus and the middle frontal gyrus, were subdivided into 2 or 3 regions while most small ROIs, such as the transverse frontal gyrus and pars opercularis, were subdivided into 2 regions or not divided at all (Tables A3 & A4 in Appendix). Silhouette scores consistently dropped after 2 or 3 subdivisions. Using this approach to subparcellation resulted in 65 cortical ROIs per hemisphere in the USCBrain atlas.

Figures 9 and 10 show quantitative evaluations of a number of atlases with respect to homogeneity within parcels of rfMRI data and consistency with a set of Brodmann areas. As should be expected, as the number of parcels increases across this set of atlases, both metrics increase. This is because rfMRI data in smaller regions will tend to be more homogeneous than in larger regions. Similarly, smaller parcels can be grouped more precisely than larger ones to better match larger Brodmann areas. However, even given these observations, it is interesting that while there is some variability in relative performance at a matched number of parcels, there are no serious outliers with respect to homogeneity. Some of the coarser parcellations based entirely on functional data (Yeo-51, Yeo0114, and Power) show poorer consistency than similar parcellation densities in anatomically-based and hybrid atlases. Interestingly, all three Schaefer atlases perform well with respect to the Brodmann metric, even though they are based on parcellation of rfMRI data only.

The BCI-DNI atlas shows very similar behavior in both metrics to the Desikan atlas. This result is unsurprising because both are based on identification of gyri, although the used 40 subjects rather than the single subject in the BCI-DNI atlas. The finer parcellation in the USCBrain atlas shows somewhat better consistency with Brodmann areas but lower functional homogeneity relative to the functional-Schaeffer-100 atlas. The reverse is the case when comparing USCBrain with the anatomical Destrieux atlas, In this case, the homogeneities are approximately equal but with 150 vs 130 cortical parcels for Destrieux vs USCBrain.

### 4.3. Intersubject Variability

Subparcellations based on repeated within-subject rfMRI scans were compared to those obtained by mapping the USCBrain atlas with gyral subdivisions to the individual subject in Figures 7 and 8. Perhaps unsurprisingly, within-subject rfMRI tended to be more consistent. However, as noted above, rfMRI data are frequently unavailable and the USCBrain atlas-based parcellations can serve as a feasible alternative to obtain functional ROIs. The observation that the ARIs for the majority of ROIs fall within average interquartile range when compared to repeated functional parcellations indicates that they represent meaningful subdivisions of these gyri. This is further supported by the agreement maps shown in Figure 6 where it is clear that for the great majority of ROIs, individual differences in functional parcellations occur close to the boundaries of the subdivisions. This variability tended to be limited to a few millimeters in most cases and reflected the typical variability we see in brain anatomy across the human population. Furthermore, a single atlas, regardless of the data used, cannot capture individual variability in functional boundaries. For that purpose, an individualized parcellation is required as described for example in (Chong et al., 2017; Wang et al., 2015). Given the absence of anatomical landmarks to guide gyral subdivisions, use of the rfMRI based subdivision here serves as a workable surrogate. The plots in Figure 7, the images in Figure 8, and the probabilistic maps provided with the atlas provide the user with an indication of the degree to which these subdivisions reflect preservation of distinct functional areas within gyri.

The bilateral insula, cingulate gyrus, and post-central gyrus had very little intersubject variability. The insula’s anterior and posterior regions are divided by its principal sulcus, which can be easily identified (Bauernfeind et al., 2013; Damasio, 2005), and have very distinct functional properties. Similarly, the cingulate gyrus’s three subdivisions have been well described in the literature and have been shown to have functionally distinct properties (Vogt et al., 1987). Conversely, bilateral superior parietal gyrus, right supramarginal gyrus, left middle frontal gyrus, and left superior occipital gyrus had large confidence intervals and low medians for the ARIs showing lower repeatability in Figure 7. Regions such as the pars triangularis, angular gyrus, and temporal lobe also tended to have less agreement across the 40 subjects in their functional boundaries in Figure 6. This lack of agreement can be explained by normal variance observed across subjects in these regions. The pars triangularis is determined by two sulcal branches that extend from the Sylvian sulcus, which can be hard to identify and, in some cases, only a single branch may be present in an individual (Berge et al., 2004; Damasio, 2005; Keller et al., 2007). The superior parietal gyrus extends into the angular gyrus which connects into the occipital lobe. This region is a multisensory area that has very complex connection patterns and functional properties. Finally, the temporal lobe’s sulcal patterns have been noted to have some of the highest complexities and variability across subjects (Ono et al., 1990). We expected to see less consistency in the frontal lobe due to the presence of susceptibility-related field distortion in fMRI data. However, this does not seem to be the case, reflecting the high quality of fMRI data and preprocessing in the HCP dataset.

Users should be cautious when interpreting functional data in regions with high variability across the population and with lower consistency across sessions as they may lead to studies with lower reproducibility. We provide the confidence map in Figure 6 and ARI values in Figure 7 to give users quantitative measures of uncertainty for each region in the USCBrain atlas. The agreement probability map shown in Figure 6 can be transferred to the subject using the one-to-one correspondence established by registration. This probability map in conjunction with the labels gives probabilistic labeling of subjects where the transferred agreement map indicates the reliability of the subdivisions. This agreement probability map is packaged with the USCBrain atlas.

One other important issue to consider is interdigitation of the brain regions and networks as noted in (Braga and Buckner, 2017). They show that some organizational features of brain networks present in the individual are blurred or lost when boundaries are averaged over a group of subjects, due to interdigitated boundaries and networks. The low probability regions in the probability maps shown in Figure 6 could be an indication of interdigitation of boundaries of subparcellations.

Anatomical registration based on T1 images is, at current, the most commonly used and well-understood registration method. While parcellation methods for many atlases, including the gyral subdivisions described here, are based on functional data it is important to realize that the registration methods themselves are (in most cases) driven by anatomical rather than functional features and landmarks. For this reason, it is important to assess limitations of anatomically-driven coregistration to functionally defined boundaries, as we do in Sections 3.2 and 3.3. Given the confound between functionally-defined atlases and anatomically-driven registration, reference atlases with manually labeled and identified anatomical segmentations remain an important benchmark that we should continue to update alongside advancing techniques based on multimodal data.

### 4.4 Atlas Usage

The BCI-DNI atlas and the USCBrain atlas are intended to be used for coregistration and segmentation of individual T1W brain images. Brain labeling or coregistration is a necessary preprocessing step for many studies examining group comparisons, correlations, and regional brain analysis. Individual T1-weighted images can be used to register images of other contrast (e.g. fMRI, DTI, etc.) from the same individual using simpler rigid registration methods. Labels, as well as the agreement probability maps, can be therefore transferred from the atlas to any of the subject images to identify anatomical and functional regions. Alternatively, images can be transformed to the atlas space for voxel-wise analysis or vertex-wise analysis on the surface.

These atlases were designed for use with the BrainSuite software package (http://brainsuite.org) (Joshi et al., 2007; Shattuck and Leahy, 2002). BrainSuite has a robust registration algorithm that uses anatomical information from both the surface and volume of the brain images for accurate automated co-registration, which allows consistent surface and volume mapping to a labeled atlas. It uses a multi-step registration and refinement process based on morphological and image intensity features and known variations in human brain anatomy and is consistent with use of a detailed single-subject atlas. BrainSuite processes images at native resolution and produces tesselated cortical surfaces with a one-to-one correspondence between brain surfaces and volumes. The BCI-DNI and USCBrain atlases with anatomical and functional segmentations are packaged with the BrainSuite software (brainsuite.org/download). Compatibility with Freesurfer and FSL is provided as described in Section 2.4 and compatible files and scripts are available (http://brainsuite.org/using-atlases/). The BrainSuite processing pipeline will output surface and volumetric labels for individual subjects. The Freesurfer pipeline outputs surface labeling and FSL outputs volumetric labeling of T1W images. As seen in Figure 11, BrainSuite and Freesurfer yield very similar results. BrainSuite and FSL produce similar results subcortically in volumetric labeling, however, FSL tends to have more bleeding across the gyral boundaries.

**Figure 11:**
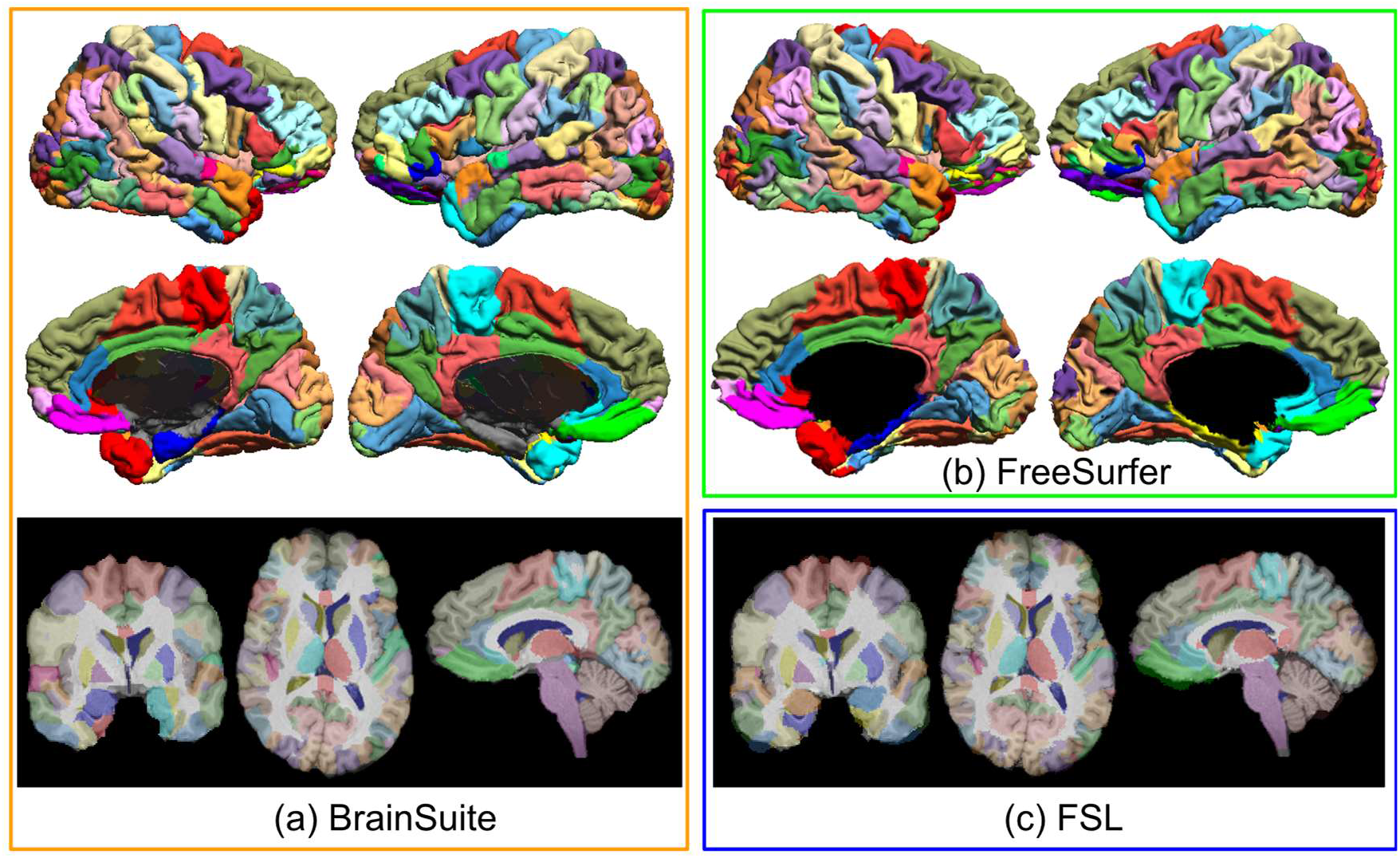
The atlas was used for labelling a representative subject using BrainSuite, FreeSurfer and FSL: (a) BrainSuite software can be used with the USCBrain atlas for surface and volume labelling; (b) FreeSurfer can be used for surface labelling while FSL (FLIRT+FNIRT) can be used for volumetric labelling.

## 5. Conclusion

In this study, we generated the anatomical BCI-DNI atlas based on single subject anatomy that was then used as a starting point for generating the hybrid USCBrain atlas with labels based on single-subject anatomy as well as functional data from a population of 40 subjects. The parcellations defined in the atlas are based on both the known anatomical landmarks defined by gyri as well as functional subdivisions of these gyri. The intended use of this atlas is to sub-parcellate cortical gyri into finer sub-divisions in the absence of anatomical landmarks in applications such as neurosurgery and studies in cognitive neuroscience. The relatively high degree of intersubject labeling agreement in the validation study indicates the utility of this atlas for labeling subjects using an anatomically driven coregistration. This atlas can be downloaded and used with our BrainSuite software (http://brainsuite.org/atlases). The source code for making the atlas is available at (https://github.com/ajoshiusc/USCBrain_atlas.git). The atlas is also compatible with FreeSurfer and FSL.

## Acknowledgments

This work is supported by the following grants: W81XWH-18-1-061, R01 NS074980, R01 NS089212, F31NS106828, K23 HD099309, and T32 MH 111360.

Data were provided by the Human Connectome Project, WU-Minn Consortium (Principal Investigators: David Van Essen and Kamil Ugurbil; 1U54MH091657) funded by the 16 NIH Institutes and Centers that support the NIH Blueprint for Neuroscience Research; and by the McDonnell Center for Systems Neuroscience at Washington University.

## Declarations of Competing Interest

none

## Appendix A. ROIs and sulci labeled in BCI-DNI atlas

**Table A1.** 95 Regions of interests (ROI) labeled on the BCI-DNI anatomical brain atlas. 66 of these regions are cortical ROIs and are labeled on the surface. *indicates non-cortical ROIs which are not labeled on the surface.

|  |  |  |
| --- | --- | --- |
| **Frontal Lobe** |  |  |
| superior frontal gyrus | middle frontal gyrus | pars opercularis |
| pars triangularis | pars orbitalis | pre-central gyrus |
| transverse frontal gyrus | gyrus rectus | middle orbitofrontal gyrus |
| anterior orbitofrontal gyrus | posterior orbitofrontal gyrus | lateral orbital gyrus |
| paracentral lobule | cingulate gyrus | subcallosal gyrus |
| **Parietal Lobe** |  |  |
| post-central gyrus | supramarginal gyrus | angular gyrus |
| superior parietal gyrus | pre-cuneus |  |
| **Temporal Lobe** |  |  |
| temporal pole | superior temporal gyrus | transverse temporal gyrus |
| middle temporal gyrus | inferior temporal gyrus | fusiform gyrus |
| parahippocampal gyrus | hippocampus* | amygdala* |
| **Occipital Lobe** |  |  |
| superior occipital gyrus | middle occipital gyrus | inferior occipital gyrus |
| lingual gyrus | cuneus |  |
| **Subcortical and Others** |  |  |
| caudate nucleus* | putamen* | globus pallidus* |
| nucleus accumbens* | thalamus* | inferior colliculus* |
| superior colliculus* | mamillary body* | pineal gland* |
| lateral ventricles* | third ventricle* | fourth ventricle* |
| cerebral aqueduct* | brainstem* | corpus callosum* |
| cerebellum* | insula |  |

**Table A2.** 26 sulci labeled on each hemisphere of the BCI-DNI anatomical brain atlas according to the BrainSuite curve protocol (http://neuroimage.usc.edu/CurveProtocol.html). Additional sulci are marked on the second set of curves totaling 39 sulci on the left hemisphere and 37 sulci on the right hemisphere. *indicates sulci as described in the original BrainSuite curve protocol.

|  |  |
| --- | --- |
| central sulcus* | precentral sulcus* |
| superior frontal sulcus* | inferior frontal sulcus* |
| ascending branch of the Sylvian fissure* | horizontal branch of the Sylvian fissure* |
| diagonal sulcus | lateral orbital sulcus* |
| frontomarginal sulcus* | cingulate sulcus* |
| paracentral sulcus* | superior supra orbital sulcus* |
| inferior supra orbital sulcus | olfactory or medial orbital sulcus* |
| H shaped sulci, mesial | H shaped sulci, lateral |
| H shaped sulci, transverse | sylvian fissure terminal split * |
| superior temporal sulcus* | inferior temporal sulcus* |
| occipito temporal sulcus* | collateral or medial occipito temporal sulcus* |
| transverse temporal sulcus* | circular sulcus* |
| postcentral sulcus* | intraparietal sulcus* |
| primary sulcus of Jensen | secondary sulcus of Jensen |
| parieto occipital sulcus* | subparietal sulcus* |
| calcarine sulcus* | calcarine sulcus terminal T |
| transverse occipital sulcus* | superior lateral occipital sulcus |
| inferior lateral occipital sulcus | anterior occipital sulcus |

## Appendix B. Silhouette Coefficients

**Table A3:** Silhouette coefficients for a different number of clusters for each anatomical ROI in the left hemisphere.

| ROI (Left hemisphere) | Average Silhouette Score |  |  |  |  |
| --- | --- | --- | --- | --- | --- |
|  | 2 Clusters | 3 Clusters | 4 Clusters | 5 Clusters | 6 Clusters |
| Precuneus | **0.46** | 0.32 | 0.25 | 0.253 | 0.26 |
| Angular gyrus | 0.54 | **0.42** | 0.36 | 0.33 | 0.30 |
| Anterior orbito-frontal gyrus | 0.21 | 0.20 | 0.23 | 0.21 | 0.24 |
| Cingulate | 0.38 | **0.46** | 0.36 | 0.33 | 0.28 |
| Cuneus | **0.30** | 0.29 | 0.28 | 0.27 | 0.24 |
| Fusiform gyrus | **0.49** | 0.27 | 0.30 | 0.21 | 0.22 |
| Gyrus Rectus | 0.41 | 0.28 | 0.22 | 0.23 | 0.15 |
| Inferior Occipital Gyrus | 0.42 | **0.47** | 0.42 | 0.42 | 0.40 |
| Inferior Temporal Gyrus | 0.51 | **0.38** | 0.31 | 0.30 | 0.24 |
| Lateral Orbito-frontal Gyrus | **0.48** | 0.31 | 0.27 | 0.22 | 0.26 |
| Lingual gyrus | **0.43** | 0.38 | 0.27 | 0.28 | 0.25 |
| Middle frontal gyrus | **0.39** | 0.33 | 0.29 | 0.24 | 0.24 |
| Middle occipital gyrus | 0.55 | **0.51** | 0.45 | 0.31 | 0.26 |
| Middle orbito-frontal gyrus | 0.15 | 0.14 | 0.13 | 0.15 | 0.17 |
| Middle temporal gyrus | 0.44 | 0.31 | **0.40** | 0.32 | 0.28 |
| Parahippocampal gyrus | 0.21 | 0.17 | 0.18 | 0.15 | 0.16 |
| Pars opercularis | **0.37** | 0.31 | 0.23 | 0.23 | 0.18 |
| Pars orbitalis | 0.29 | 0.25 | 0.23 | 0.25 | 0.28 |
| Pars triangularis | 0.29 | **0.34** | 0.33 | 0.31 | 0.29 |
| Post-central gyrus | **0.41** | 0.38 | 0.29 | 0.32 | 0.30 |
| Posterior orbito-frontal gyrus | 0.15 | 0.18 | 0.21 | 0.22 | 0.24 |
| Pre-central gyrus | **0.39** | 0.30 | 0.33 | 0.29 | 0.27 |
| Subcallosal gyrus | 0.13 | 0.16 | 0.24 | 0.25 | 0.25 |
| Superior frontal gyrus | **0.45** | 0.26 | 0.21 | 0.23 | 0.27 |
| Superior occipital gyrus | **0.41** | 0.33 | 0.31 | 0.27 | 0.22 |
| Superior parietal gyrus | **0.33** | 0.33 | 0.29 | 0.33 | 0.33 |
| Supramarginal gyrus | **0.48** | 0.39 | 0.35 | 0.32 | 0.31 |
| Superior temporal gyrus | 0.31 | **0.28** | 0.27 | 0.31 | 0.28 |
| Temporal pole | 0.22 | 0.21 | 0.20 | 0.19 | 0.17 |
| Transverse frontal gyrus | **0.38** | 0.25 | 0.24 | 0.21 | 0.22 |
| Transverse temporal gyrus | 0.23 | 0.17 | 0.15 | 0.19 | 0.19 |
| Insula | **0.37** | 0.28 | 0.21 | 0.18 | 0.20 |

**Table A4:** Silhouette coefficients for a different number of clusters for each anatomical ROI in the left hemisphere.

| ROI (Right hemisphere) | Average Silhouette Score |  |  |  |  |
| --- | --- | --- | --- | --- | --- |
|  | 2<br>Clusters | 3<br>Clusters | 4<br>Clusters | 5<br>Clusters | 6<br>Clusters |
| Precuneus | **0.48** | 0.35 | 0.22 | 0.29 | 0.25 |
| Angular gyrus | 0.28 | **0.37** | 0.33 | 0.31 | 0.33 |
| Anterior orbito-frontal gyrus | 0.21 | 0.18 | 0.19 | 0.20 | 0.21 |
| Cingulate | 0.34 | **0.43** | 0.37 | 0.31 | 0.24 |
| Cuneus | **0.36** | 0.28 | 0.28 | 0.28 | 0.25 |
| Fusiforme gyrus | **0.49** | 0.29 | 0.24 | 0.21 | 0.16 |
| Gyrus Rectus | 0.21 | 0.14 | 0.15 | 0.15 | 0.11 |
| Inferior Occipital Gyrus | 0.34 | **0.36** | 0.38 | 0.39 | 0.33 |
| Inferior Temporal Gyrus | 0.58 | **0.37** | 0.31 | 0.25 | 0.22 |
| Lateral Orbito-frontal Gyrus | **0.29** | 0.16 | 0.20 | 0.18 | 0.17 |
| Lingual gyrus | **0.34** | 0.20 | 0.28 | 0.26 | 0.25 |
| Middle frontal gyrus | **0.37** | 0.27 | 0.27 | 0.23 | 0.23 |
| Middle occipital gyrus | 0.40 | **0.51** | 0.36 | 0.36 | 0.32 |
| Middle orbito-frontal gyrus | 0.16 | 0.15 | 0.15 | 0.15 | 0.15 |
| Middle temporal gyrus | 0.36 | 0.33 | **0.38** | 0.32 | 0.26 |
| Parahippocampal gyrus | 0.26 | 0.15 | 0.13 | 0.17 | 0.15 |
| Pars opercularis | **0.38** | 0.26 | 0.23 | 0.19 | 0.14 |
| Pars orbitalis | 0.22 | 0.20 | 0.21 | 0.22 | 0.22 |
| Pars trianularis | 0.26 | **0.24** | 0.17 | 0.20 | 0.21 |
| Post-central gyrus | **0.43** | 0.39 | 0.37 | 0.31 | 0.27 |
| Posterior orbito-frontal gyrus | 0.13 | 0.15 | 0.16 | 0.16 | 0.18 |
| Pre-central gyrus | **0.36** | 0.32 | 0.37 | 0.33 | 0.29 |
| Subcallosal gyrus | 0.17 | 0.19 | 0.23 | 0.27 | 0.31 |
| Superior frontal gyrus | **0.46** | 0.26 | 0.23 | 0.23 | 0.28 |
| Superior occipital gyrus | **0.57** | 0.45 | 0.33 | 0.23 | 0.27 |
| Superior parietal gyrus | **0.37** | 0.33 | 0.32 | 0.29 | 0.28 |
| Supramarginal gyrus | **0.39** | 0.34 | 0.32 | 0.33 | 0.35 |
| Superior temporal gyrus | 0.35 | **0.36** | 0.27 | 0.24 | 0.26 |
| Temporal pole | 0.19 | 0.19 | 0.19 | 0.18 | 0.15 |
| Transvers frontal gyrus | **0.35** | 0.22 | 0.19 | 0.19 | 0.19 |
| Transverse temporal gyrus | 0.20 | 0.14 | 0.15 | 0.14 | 0.17 |
| Insula | **0.30** | 0.23 | 0.20 | 0.19 | 0.14 |

## Notes

http://brainsuite.org/uscbrainatlas

### Competing Interest Statement

The authors have declared no competing interest.

## REFERENCES

Andersson, J.L., Jenkinson, M., Smith, S., 2007. Non-linear registration, aka spatial normalisation. FMRIB technial report TR07JA2 22.

Arslan, S., Ktena, S.I., Makropoulos, A., Robinson, E.C., Rueckert, D., Parisot, S., 2017. Human brain mapping: A systematic comparison of parcellation methods for the human cerebral cortex. NeuroImage. https://doi.org/10.1016/j.neuroimage.2017.04.014

Bauernfeind, A.L., de Sousa, A.A., Avasthi, T., Dobson, S.D., Raghanti, M.A., Lewandowski, A.H., Zilles, K., Semendeferi, K., Allman, J.M., Craig, A.D. (Bud), Hof, P.R., Sherwood, C.C., 2013. A volumetric comparison of the insular cortex and its subregions in primates. Journal of Human Evolution 64, 263–279. https://doi.org/10.1016/j.jhevol.2012.12.003

Behrens, T.E.J., Johansen-Berg, H., Woolrich, M.W., Smith, S.M., Wheeler-Kingshott, C.A.M., Boulby, P.A., Barker, G.J., Sillery, E.L., Sheehan, K., Ciccarelli, O., Thompson, A.J., Brady, J.M., Matthews, P.M., 2003. Non-invasive mapping of connections between human thalamus and cortex using diffusion imaging. Nat Neurosci 6, 750–757. https://doi.org/10.1038/nn1075

Berge, J., Vader, W., Lockhart, S., 2004. A survey of amphipod associates of sea urchins, with description of new species in the genera Lepidepecreella (Lysianassoidea: lepidepecreellid group) and Notopoma (Photoidea: Ischyroceridae) from Antarctic cidarids. Stuttgart, New York:Thieme. https://doi.org/10.1016/j.dsr2.2004.06.031

Bhushan, C., Chong, M., Choi, S., Joshi, A.A., Haldar, J.P., 2016. Temporal Non-Local Means Filtering Reveals Real-Time Whole-Brain Cortical Interactions in Resting fMRI. PloS one 11, 1–22. https://doi.org/10.1371/journal.pone.0158504

Blumensath, T., Behrens, T.E.J., Smith, S.M., 2012. Resting-State FMRI Single Subject Cortical Parcellation Based on Region Growing, in: International Conference on Medical Image Computing and Computer-Assisted Intervention. Springer, pp. 188–195. https://doi.org/10.1007/978-3-642-33418-4_24

Braga, R.M., Buckner, R.L., 2017. Parallel Interdigitated Distributed Networks within the Individual Estimated by Intrinsic Functional Connectivity. Neuron 95, 457-471.e5. https://doi.org/10.1016/j.neuron.2017.06.038

Calhoun, V.D., Liu, J., Adali, T., 2009. A review of group ICA for fMRI data and ICA for joint inference of imaging, genetic, and ERP data. NeuroImage 45, S163—-S172. https://doi.org/10.1016/j.neuroimage.2008.10.057

Chakravarty, M.M., Bertrand, G., Hodge, C.P., Sadikot, A.F., Collins, D.L., 2006. The creation of a brain atlas for image guided neurosurgery using serial histological data. NeuroImage 30, 359–376. https://doi.org/10.1016/j.neuroimage.2005.09.041

Chong, M., Bhushan, C., Joshi, A.A., Choi, S., Haldar, J.P., Shattuck, D.W., Spreng, R.N., Leahy, R.M., 2017. Individual parcellation of resting fMRI with a group functional connectivity prior. NeuroImage 156, 87–100. https://doi.org/10.1016/j.neuroimage.2017.04.054

Cointepas, Y., Mangin, J.-F., Garnero, L., Poline, J.-B., Benali, H., 2001. BrainVISA: software platform for visualization and analysis of multi-modality brain data. Neuroimage 13, 98.

Cordes, D., Haughton, V., Carew, J.D., Arfanakis, K., Maravilla, K., 2002. Hierarchical clustering to measure connectivity in fMRI resting-state data. Magnetic Resonance Imaging 20, 305–317. https://doi.org/10.1016/S0730-725X(02)00503-9

Craddock, R.C., James, G.A., Iii, P.E.H., Hu, X.P., Mayberg, H.S., 2013. A whole brain fMRI atlas spatial Generated via Spatially Constrained Spectral Clustering_ Craddock, James 2011 .pdf. Human brain mapping 33, 1914–1928. https://doi.org/10.1002/hbm.21333.A

Damasio, H., 2005. Human brain anatomy in computerized images. Oxford University Press, USA.

de Amorim, R.C., Hennig, C., 2015. Recovering the number of clusters in data sets with noise features using feature rescaling factors. Information Sciences 324, 126–145. https://doi.org/10.1016/j.ins.2015.06.039

Desbrun, M., Polthier, K., 2007. Discrete Differential Geometry. Computer Aided Geometric Design 24, 427. https://doi.org/10.1016/j.cagd.2007.07.005

Desikan, R.S., Ségonne, F., Fischl, B., Quinn, B.T., Dickerson, B.C., Blacker, D., Buckner, R.L., Dale, A.M., Maguire, R.P., Hyman, B.T., others, 2006. An automated labeling system for subdividing the human cerebral cortex on MRI scans into gyral based regions of interest. Neuroimage 31, 968–980.

Dickie, D.A., Shenkin, S.D., Anblagan, D., Lee, J., Blesa Cabez, M., Rodriguez, D., Boardman, J.P., Waldman, A., Job, D.E., Wardlaw, J.M., 2017. Whole Brain Magnetic Resonance Image Atlases: A Systematic Review of Existing Atlases and Caveats for Use in Population Imaging. Front. Neuroinform. 11. https://doi.org/10.3389/fninf.2017.00001

Do Carmo, M.P., 2016. Differential geometry of curves and surfaces: revised and updated second edition. Courier Dover Publications.

Duvernoy, H.M., 1999. The Human Brain. Springer Vienna, Vienna. https://doi.org/10.1007/978-3-7091-6792-2

Essen, D.C.V., Drury, H.A., 1997. Structural and Functional Analyses of Human Cerebral Cortex Using a Surface-Based Atlas. J. Neurosci. 17, 7079–7102.

Fan, L., Li, H., Zhuo, J., Zhang, Y., Wang, J., Chen, L., Yang, Z., Chu, C., Xie, S., Laird, A.R., others, 2016. The human brainnetome atlas: a new brain atlas based on connectional architecture. Cerebral Cortex 26, 3508–3526.

Fischl, B., 2012. FreeSurfer. NeuroImage 62, 774–781. https://doi.org/10.1016/j.neuroimage.2012.01.021

Fischl, B., Sereno, M.I., Dale, A.M., 1999. Cortical Surface-Based Analysis. NeuroImage 9, 195–207. https://doi.org/10.1006/nimg.1998.0396

Fischl, B., van der Kouwe, A., Destrieux, C., Halgren, E., Ségonne, F., Salat, D.H., Busa, E., Seidman, L.J., Goldstein, J., Kennedy, D., Caviness, V., Makris, N., Rosen, B., Dale, A.M., 2004. Automatically Parcellating the Human Cerebral Cortex. Cereb Cortex 14, 11–22. https://doi.org/10.1093/cercor/bhg087

Glasser, M.F., Coalson, T.S., Robinson, E.C., Hacker, C.D., Harwell, J., Yacoub, E., Ugurbil, K., Andersson, J., Beckmann, C.F., Jenkinson, M., Smith, S.M., Van Essen, D.C., 2016. A multi-modal parcellation of human cerebral cortex. Nature 536, 171–178. https://doi.org/10.1038/nature18933

Glasser, M.F., Goyal, M.S., Preuss, T.M., Raichle, M.E., Van Essen, D.C., 2014. Trends and properties of human cerebral cortex: correlations with cortical myelin content. Neuroimage 93 Pt 2, 165–175. https://doi.org/10.1016/j.neuroimage.2013.03.060

Glasser, M.F., Sotiropoulos, S.N., Wilson, J.A., Coalson, T.S., Fischl, B., Andersson, J.L., Xu, J., Jbabdi, S., Webster, M., Polimeni, J.R., Van Essen, D.C., Jenkinson, M., 2013. The minimal preprocessing pipelines for the Human Connectome Project. NeuroImage 80, 105–124. https://doi.org/10.1016/j.neuroimage.2013.04.127

Glasser, M.F., Van Essen, D.C., 2011. Mapping Human Cortical Areas In Vivo Based on Myelin Content as Revealed by T1-and T2-Weighted MRI. Journal of Neuroscience 31, 11597–11616. https://doi.org/10.1523/JNEUROSCI.2180-11.2011

Gordon, E.M., Laumann, T.O., Adeyemo, B., Huckins, J.F., Kelley, W.M., Petersen, S.E., 2016. Generation and Evaluation of a Cortical Area Parcellation from Resting-State Correlations. Cerebral Cortex 26, 288–303. https://doi.org/10.1093/cercor/bhu239

Holmes, C.J., Hoge, R., Collins, L., Woods, R., Toga, A.W., Evans, A.C., 1998. Enhancement of MR Images Using Registration for Signal Averaging. Journal of Computer Assisted Tomography 22, 324–333.

Johansen-Berg, H., Behrens, T.E.J., Robson, M.D., Drobnjak, I., Rushworth, M.F.S., Brady, J.M., Smith, S.M., Higham, D.J., Matthews, P.M., 2004. Changes in connectivity profiles define functionally distinct regions in human medial frontal cortex. Proceedings of the National Academy of Sciences 101, 13335–13340. https://doi.org/10.1073/pnas.0403743101

Joshi, A.A., Chong, M., Li, J., Choi, S., Leahy, R.M., 2018. Are you thinking what I’m thinking? Synchronization of resting fMRI time-series across subjects. NeuroImage 172, 740–752. https://doi.org/10.1016/j.neuroimage.2018.01.058

Joshi, A.A., Shattuck, D.W., Damasio, H., Leahy, R.M., 2012a. Geodesic curvature flow on surfaces for automatic sulcal delineation, in: Proceedings - International Symposium on Biomedical Imaging. IEEE, pp. 430–433. https://doi.org/10.1109/ISBI.2012.6235576

Joshi, A.A., Shattuck, D.W., Leahy, R.M., 2012b. A Method for Automated Cortical Surface Registration and Labeling, in: Dawant, B.M., Christensen, G.E., Fitzpatrick, J.M., Rueckert, D. (Eds.), Biomedical Image Registration, Lecture Notes in Computer Science. Springer Berlin Heidelberg, Berlin, Heidelberg, pp. 180–189. https://doi.org/10.1007/978-3-642-31340-0_19

Joshi, A.A., Shattuck, D.W., Thompson, P.M., Leahy, R.M., 2007. Surface-Constrained Volumetric Brain Registration Using Harmonic Mappings. IEEE Trans. Med. Imaging 26, 1657–1669.

Keller, S.S., Highley, J.R., Garcia-Finana, M., Sluming, V., Rezaie, R., Roberts, N., 2007. Sulcal variability, stereological measurement and asymmetry of Broca’s area on MR images. J Anat 211, 534–555. https://doi.org/10.1111/j.1469-7580.2007.00793.x

Kuhn, H.W., 1955. The Hungarian method for the assignment problem. Naval Research Logistics 2, 83–97. https://doi.org/10.1002/nav.3800020109

Li, J., Choi, S., Joshi, A.A., Wisnowski, J.L., Leahy, R.M., 2020. Temporal non-local means filtering for studies of intrinsic brain connectivity from individual resting fMRI. Medical Image Analysis 61, 101635. https://doi.org/10.1016/j.media.2020.101635

Li, J., Choi, S., Joshi, A.A., Wisnowski, J.L., Leahy, R.M., 2018. Global PDF-based temporal non-local means filtering reveals individual differences in brain connectivity, in: 2018 IEEE 15th International Symposium on Biomedical Imaging (ISBI 2018). pp. 15–19. https://doi.org/10.1109/ISBI.2018.8363513

Mailo, J., Tang-Wai, R., 2015. Insight into the precuneus: a novel seizure semiology in a child with epilepsy arising from the right posterior precuneus. Epileptic Disord 17, 321–327. https://doi.org/10.1684/epd.2015.0759

Makris, N., Meyer, J.W., Bates, J.F., Yeterian, E.H., Kennedy, D.N., Caviness, V.S., 1999. MRI-Based Topographic Parcellation of Human Cerebral White Matter and Nuclei. NeuroImage 9, 18–45. https://doi.org/10.1006/nimg.1998.0384

Margulies, D.S., Vincent, J.L., Kelly, C., Lohmann, G., Uddin, L.Q., Biswal, B.B., Villringer, A., Castellanos, F.X., Milham, M.P., Petrides, M., 2009. Precuneus shares intrinsic functional architecture in humans and monkeys. PNAS 106, 20069–20074. https://doi.org/10.1073/pnas.0905314106

Mazziotta, J.C., Toga, A.W., Evans, A., Fox, P., Lancaster, J., 1995. A Probabilistic Atlas of the Human Brain: Theory and Rationale for Its Development. NeuroImage 2, 89–101. https://doi.org/10.1006/nimg.1995.1012

Miller, M.B., Donovan, C.-L., Van Horn, J.D., German, E., Sokol-Hessner, P., Wolford, G.L., 2009. Unique and persistent individual patterns of brain activity across different memory retrieval tasks. NeuroImage 48, 625–635. https://doi.org/10.1016/j.neuroimage.2009.06.033

Mori, S., Oishi, K., Faria, A.V., Miller, M.I., 2013. Atlas-Based Neuroinformatics via MRI: Harnessing Information from Past Clinical Cases and Quantitative Image Analysis for Patient Care. Annu. Rev. Biomed. Eng. 15, 71–92. https://doi.org/10.1146/annurev-bioeng-071812-152335

Ono, M., Kubik, S., Abernathey, C.D., 1990. Atlas of the Cerebral Sulci. Stuttgart, New York:Thieme.

Pantazis, D., Joshi, A., Jiang, J., Shattuck, D.W., Bernstein, L.E., Damasio, H., Leahy, R.M., 2010. Comparison of landmark-based and automatic methods for cortical surface registration. Neuroimage 49, 2479–2493.

Power, J.D., Cohen, A.L., Nelson, S.M., Wig, G.S., Barnes, K.A., Church, J.A., Vogel, A.C., Laumann, T.O., Miezin, F.M., Schlaggar, B.L., Petersen, S.E., 2011. Functional Network Organization of the Human Brain. Neuron 72, 665–678. https://doi.org/10.1016/j.neuron.2011.09.006

Rand, W.M., 1971. Objective Criteria for the Evaluation of Clustering Methods. Journal of the American Statistical Association 66, 846–850. https://doi.org/10.2307/2284239

Rokach, L., Maimon, O., 2005. Clustering Methods, in: Maimon, O., Rokach, L. (Eds.), Data Mining and Knowledge Discovery Handbook. Springer-Verlag, New York, pp. 321–352. https://doi.org/10.1007/0-387-25465-X_15

Rolls, E.T., Huang, C.-C., Lin, C.-P., Feng, J., Joliot, M., 2020. Automated anatomical labelling atlas 3. NeuroImage 206, 116189. https://doi.org/10.1016/j.neuroimage.2019.116189

Rousseeuw, P.J., 1987. Rousseeuw, P.J.: Silhouettes: A Graphical Aid to the Interpretation and Validation of Cluster Analysis. Comput. Appl. Math. 20, 53-65. Journal of Computational & Applied Mathematics 20, 53–65.

Schaefer, A., Kong, R., Gordon, E.M., Laumann, T.O., Zuo, X.-N., Holmes, A.J., Eickhoff, S.B., Yeo, B.T.T., 2018. Local-Global Parcellation of the Human Cerebral Cortex from Intrinsic Functional Connectivity MRI. Cerebral Cortex 28, 3095–3114. https://doi.org/10.1093/cercor/bhx179

Shattuck, D.W., Leahy, R.M., 2002. BrainSuite: An Automated Cortical Surface Identification Tool. Medical Image Analysis 8, 129–142.

Shi, J., Malik, J., 2005. Normalized Cuts and Image Segmentation Normalized Cuts and Image Segmentation. IEEE Transactions on pattern analysis and machine intelligence 22, 888–905. https://doi.org/10.1109/CVPR.1997.609407

Taylor, K.N., Joshi, A.A., Hirfanoglu, T., Grinenko, O., Liu, P., Wang, X., Gonzalez-Martinez, J.A., Leahy, R.M., Mosher, J.C., Nair, D.R., 2021. Validation of semi-automated anatomically labeled SEEG contacts in a brain atlas for mapping connectivity in focal epilepsy. Epilepsia Open 6, 493–503. https://doi.org/10.1002/epi4.12499

Taylor, K.N., Joshi, A.A., Li, J., Gonzalez-Martinez, J.A., Wang, X., Leahy, R.M., Nair, D.R., Mosher, J.C., 2020. The FAST graph: A novel framework for the anatomically-guided visualization and analysis of cortico-cortical evoked potentials. Epilepsy Research 161, 106264. https://doi.org/10.1016/j.eplepsyres.2020.106264

Van Essen, D.C., Smith, S.M., Barch, D.M., Behrens, T.E.J., Yacoub, E., Ugurbil, K., WU-Minn HCP Consortium, 2013. The WU-Minn Human Connectome Project: an overview. Neuroimage 80, 62– 79. https://doi.org/10.1016/j.neuroimage.2013.05.041

Vogt, B.A., Pandya, D.N., Rosene, D.L., 1987. Cingulate cortex of the rhesus monkey: I. Cytoarchitecture and thalamic afferents. J. Comp. Neurol. 262, 256–270. https://doi.org/10.1002/cne.902620207

Wang, D., Buckner, R.L., Fox, M.D., Holt, D.J., Holmes, A.J., Stoecklein, S., Langs, G., Pan, R., Qian, T., Li, K., Baker, J.T., Stufflebeam, S.M., Wang, K., Wang, X., Hong, B., Liu, H., 2015. Parcellating cortical functional networks in individuals. Nat Neurosci 18, 1853–1860. https://doi.org/10.1038/nn.4164

Yeo, B.T.T., Krienen, F.M., Sepulcre, J., Sabuncu, M.R., Lashkari, D., Hollinshead, M., Roffman, J.L., Smoller, J.W., Zöllei, L., Polimeni, J.R., Fischl, B., Liu, H., Buckner, R.L., 2011. The organization of the human cerebral cortex estimated by intrinsic functional connectivity. J. Neurophysiol. 106, 1125–1165. https://doi.org/10.1152/jn.00338.2011

Zhi, D., King, M., Diedrichsen, J., 2021. Evaluating brain parcellations using the distance controlled boundary coefficient (preprint). Neuroscience. https://doi.org/10.1101/2021.05.11.443151

Zilles, K., Amunts, K., 2010. Centenary of Brodmann’s map — conception and fate. Nat Rev Neurosci 11, 139–145. https://doi.org/10.1038/nrn2776

